# PP2A/B55α substrate recruitment as defined by the retinoblastoma-related protein p107

**DOI:** 10.1101/2021.03.02.433577

**Authors:** Holly Fowle, Ziran Zhao, Qifang Xu, Xinru Wang, Mary Adeyemi, Felicity Feiser, Alison Kurimchak, Arminja N. Kettenbach, Rebecca Page, Wolfgang Peti, Roland L. Dunbrack, Xavier Graña

## Abstract

Protein phosphorylation is a reversible post-translation modification essential in cell signaling. This study addresses a long-standing question as to how the most abundant serine/threonine Protein Phosphatase 2 (PP2A) holoenzyme, PP2A/B55α, specifically recognizes substrates and presents them to the enzyme active site. Here, we show how the PP2A regulatory subunit B55α recruits p107, a pRB-related tumor suppressor and B55α substrate. Using molecular and cellular approaches, we identified a conserved region 1 (R1, residues 615-626) encompassing the strongest p107 binding site. This enabled us to identify an “HxRVxxV^619-625^” short linear motif (SLiM) in p107 as necessary for B55α binding and dephosphorylation of the proximal pSer-615 *in vitro* and in cells. Numerous B55α/PP2A substrates, including TAU, contain a related SLiM C-terminal from a proximal phosphosite, “**p**[**ST**]-**P-**x(5-10)-[**RK**]-**V**-x-x-[**VI**]-**R**”. Mutation of conserved SLiM residues in TAU dramatically inhibits dephosphorylation by PP2A/B55α, validating its generality. A data-guided computational model details the interaction of residues from the conserved p107 SLiM, the B55α groove, and phosphosite presentation. Altogether these data provide key insights into PP2A/B55α mechanisms of substrate recruitment and active site engagement, and also facilitate identification and validation of new substrates, a key step towards understanding PP2A/B55〈 role in multiple cellular processes.

## Introduction

Protein phosphorylation is a reversible post-translational modification that is critical for the regulation of signaling and other cellular processes. It is estimated that a third of all cellular proteins are phosphorylated (Ficarro *et al*, 2002), with more than 98% of those phosphorylation events occurring on serine and threonine residues (Olsen *et al*, 2006). The opposing processes of phosphorylation and dephosphorylation are catalyzed by protein kinases and phosphatases, respectively. Despite the fundamental importance of dephosphorylation for normal cell physiology, the mechanisms of substrate recognition by protein phosphatases are only poorly understood (reviewed in Brautigan & Shenolikar, 2018).

Members of the <u>p</u>hospho<u>p</u>rotein <u>p</u>hosphatase (PPP) family of serine/threonine phosphatases are responsible for the majority of dephosphorylation in eukaryotic cells, with protein phosphatase 1 (PP1) and protein phosphatase 2A (PP2A) accounting for more than 90% of the total phosphatase activity (Moorhead *et al*, 2007; Virshup & Shenolikar, 2009). Within the PPP family, the molecular details of substrate recognition have been best studied for PP1 and calcineurin (or PP2B), and involve the recognition of defined short linear motifs or SLiMs including RVxF, ϕϕ (where ϕ refers to a hydrophobic residue), SILK, among others for PP1 (Choy *et al*, 2014; Kumar *et al*, 2018); and LxVP and PxIxIT for PP2B. These motifs are characterized by the presence of three or four core interacting amino acids that are part of a 4-10 amino acid stretch within intrinsically disordered regions of regulator and/or substrate proteins. This SLiM-mediated specific targeting of substrates or regulatory proteins to PPPs is essential for the temporal and spatial coordination of its functions (reviewed in Brautigan & Shenolikar, 2018; Heroes *et al*, 2013).

PP2A is a highly conserved Ser/Thr protein phosphatase that makes up close to 1% of total cellular protein in some tissue types, making it one of the most abundant enzymes (Fowle *et al*, 2019). As a multimeric protein, PP2A can exist as a heterodimeric “core enzyme”, consisting of a scaffold (A) subunit and a catalytic (C) subunit, or a heterotrimeric holoenzyme, in which the core dimer complexes with a B subunit (reviewed in Shi, 2009). The B subunit of PP2A can be subdivided into one of four major families (B55, B56, B72, and B93), each consisting of two to five isoforms and numerous splice variants. Apart from its role in subcellular localization, the B subunits are thought to be the key determinant of substrate specificity for the PP2A complex (reviewed in Fowle *et al*., 2019; Kurimchak & Graña, 2012, 2015; Virshup & Shenolikar, 2009). Indeed, a PP2A substrate SLiM (LxxIxE) has recently been identified for the B56 family of PP2A regulatory subunits, which form a HEAT repeat fold. The binding affinity of LxxIxE can be modulated by phosphorylation (phosphorylation leads to tighter binding) and it binds in a binding pocket between HEAT repeats 3 and 4. Furthermore, it was recently shown that additional factors including conserved, dynamic charged:charged interaction modulate substrate specificity for B56 (Wang *et al*, 2020). This combined molecular and cellular data can be leveraged to validate known B56/PP2A substrates and, most importantly, identify potential novel B56 substrates (Hertz *et al*, 2016; Wang *et al*, 2016). In contrast, most of this information is missing for all other B family, limiting our ability to understand substrate recruitment.

B55α (4 isoforms, α, β, ɣ, δ; our work focuses on the α-isoform of B55) is ubiquitously expressed and the most abundant regulatory subunit of PP2A (Kim *et al*, 2014; Wang *et al*, 2015). Critically, all known PP2A/B55α dependent substrates have key functions in cell division, differentiation, and survival, and are found to be dysregulated in cancer and Alzheimer’s disease (reviewed in Fowle *et al*., 2019). The structure of the PP2A/B55α holoenzyme allowed for the suggestion of a mechanism for targeting the substrate Tau. Specifically, it was proposed the large negatively charged patch on B55α recruits Lys-rich domains of TAU (Xu *et al*, 2008), only to be shown that other substrates must use different molecular recognition motifs (Jayadeva *et al*, 2010). Recently, proteomics data suggested that B55α preferred substrates that are phosphorylated by Ser/Thr-Pro directed kinases (Cundell *et al*, 2016; Zhao *et al*, 2019). However, no substrate recruitment mechanism has been identified for B55α and thus further insights are of key importance to understand how substrate engage B55.

Here, by combining and leveraging molecular and cellular data, we characterize the recruitment of the PP2A/B55α specific substrate p107 (Garriga *et al*, 2004; Jayadeva *et al*., 2010; Kolupaeva *et al*, 2013; Kurimchak *et al*, 2013). p107 is a pRB-related tumor suppressor with key regulatory roles in the cell cycle (reviewed in Kurimchak & Graña, 2015). Our data leads to the identification of a conserved SLiM in key B55α substrates and enables us to develop a structural model for B55α substrate recognition that enables us to predict how other substrates bind B55α. Since modulation of PP2A activity is actively being explored in many cancer types, our data will further help in turning PP2A in a key cancer drug target.

## Results

### p107 binds to B55

p107 (Retinoblastoma-like protein 1, RBL1) is a multi-domain protein. We have previously shown that the p107 “spacer” region (p107 residues 585-780, which link the retinoblastoma conserved regions A and B) is sufficient for PP2A/B55α binding (Jayadeva *et al*., 2010). IUPred2a bioinformatics analysis (Mészáros *et al*, 2018) shows that this linker region is intrinsically disordered (Fig. 1A). Aligning the intrinsically disordered p107 linker region from 60 different species enabled the identification of three highly conserved regions (termed R1, R2, and R3) (Fig 1B, suppl. Fig 1A). Indeed, the conservation of R1 and R2 extended to a conserved family member p130 (RBL2) (Fig. 1C). Within the spacer there are also three potential CDK kinase phosphorylation sites (p107 S615, S640 and S650).

**Figure 1.**
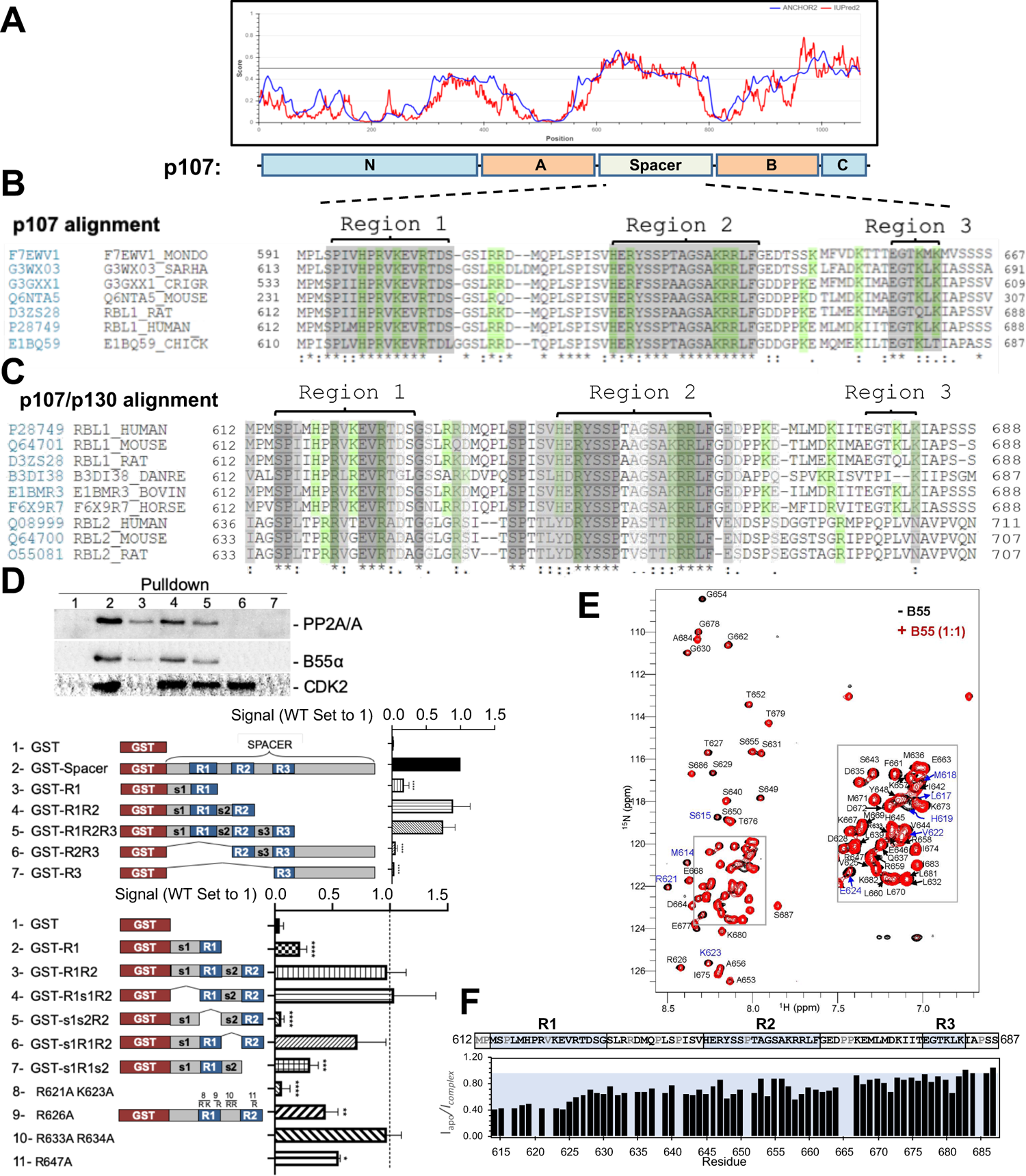
The intrinsically-disordered spacer region of p107 contains 3 highly conserved regions (R1, R2, and R3), of which R1 is required for B55α binding and R2 enhances the binding interaction as determined via mutational analysis and NMR. (A) The spacer and the C-terminus of p107 are intrinsically disordered (IUPRED server). (B) Clustal W alignment of conserved amino acid sequences of the p107 spacer from different species. Three highly conserved regions within the spacer were identified, which are highlighted in grey and named as Region I (R1), Region 2 (R2) and Region 3 (R3). Positively charged residues are highlighted green. (C) Clustal W alignment of conserved amino acid sequences of the p107 and p130 spacer from the indicated species. Conserved residues are highlighted in shades of gray. Positively charged residues are highlighted green. (D) A GST-p107 spacer construct was used as a template to systematically delete indicated regions and mutate positively charged amino acids in R1 and R2. Pull down assays with the indicated fusion proteins from U2-OS lysates were performed and binding of the indicated proteins was determined by western blot analysis. Experiments were performed in triplicate and quantification values represent the mean ± standard deviation (SD). (E) Overlay of the 2D [1H,15N] HSQC spectra of 15N-labeled p107 (M612-S687) in the presence (red) and absence (black) of purified monomeric full length B55α (M1-N447) (see methods for B55α purification). (F) p107 (M612-S687) sequence is shown above the peak intensity ratios for data shown in (E). Prolines, residues not assigned (M612 and R634), overlapping residue (V622) were labeled in grey. Residues corresponding to R1, R2, and R3 regions are highlighted in blue.

### R1 is necessary and sufficient for B55α binding

To determine the contribution of each conserved region, we generated p107 deletion mutants and performed GST pull-down assays using U-2 OS whole cell lysates. Fig. 1D shows that a mutant lacking residues C-terminal of R2 binds B55α similarly to the full construct, indicating that residues C-terminal to the R2 domain are dispensable for B55α binding. R1 alone can bind B55α, although to a slightly lesser extent than R1/R2 (lane 3), while mutants lacking regions containing R1 do not bind B55α (lanes 6 and 7). Additional experiments highlighted that R1 is the key B55α binding region and that R2 enhances the interaction, but without R1 cannot recruit B55α alone (Fig 1D, Suppl. Fig. 1B). Conversely, binding of p107 to CDK2 strictly depends on the presence of R2 (lanes 2, 4, 5, and 6), which includes an RxL motif necessary for Cyclin/CDK binding (Adams *et al*, 1996; Chen *et al*, 1996).

It has been shown that charged:charged interactions are central for B55α:TAU substrate recruitment. To test if charged:charged interactions are also important for p107 recruitment, we generated p107 R621A and K623A variants (in R1) and R647A variant (in R2) using GST-p107-R1R2. As shown in Fig. 1D and Suppl. Fig. 1B, positive residues in R1 and R2, but not the connecting “linker,” led to a reduction in binding to B55α. Thus, positively charged residues participate in binding to B55α.

We also used NMR spectroscopy to gain further atomic resolution insights into the interaction between p107 and B55α. The 2D [^1^H,^15^N] HSQC spectrum of ^15^N-labeled p107 (residues M612-S687, which include R1, R2 and R3) shows all hallmarks of an intrinsically disordered protein (IDP), with a highly limited proton chemical shift dispersion due to the lack of a hydrogen bond network in secondary structure elements. This experimentally validates the bioinformatics data. An overlay of the 2D [^1^H,^15^N] HSQC spectrum of ^15^N-labeled p107_M612-S687_ in the presence and absence of purified B55α shows that ∼15 peaks have reduced intensity, typical for an IDP:protein interaction (Fig. 1E-F). Upon completion of the sequence-specific backbone assignment of p107_M612-S687_, we identified that cross-peaks corresponding to p107 residues M614-E624 were most significantly broadened upon binding to B55α (Fig. 1F). Consistent with the mutation data, these residues form the core of R1. Furthermore, residues within the linker region and R2 also showed peak intensity attenuations, consistent with additional weaker interactions in R2 contributing and enhancing the interaction of R1. Taken together, the NMR data reinforces that the p107 R1 region mediates key contacts with B55α.

### p107 interacts with specific B55α surface residues

The recently identified LxxIxE B56 SLiM binds to a highly conserved binding pocket on B56. Thus, to identify B55α residues that mediate p107 binding, we analyzed B55α conservation. B55α adopts a β-propeller fold. The most highly conserved residues cluster at the top of the β-propeller and a fraction of them form a highly negatively charged patch (Fig. 2A-B, Suppl. Fig. 2A). Based on this analysis and the observation of a prominent groove on the surface of B55α blade 4 (Fig. 2A-B, Suppl. Fig. 2A) that might function as a potential guide for substrates to reach the catalytic site of PP2A/C, we generated 19 Myc-tagged B55α variants (either Ala substitutions or charge reversals; Fig. 2C, see Suppl. Fig. 2B for complete list). All 19 B55α variants were tested in co-immunoprecipitation assays to measure binding to p107. Furthermore, we also tested binding for two additional PP2A:B55α substrates, pRB and KSR1, to test if this B55α interaction surface is shared by different substrates.

**Figure 2.**
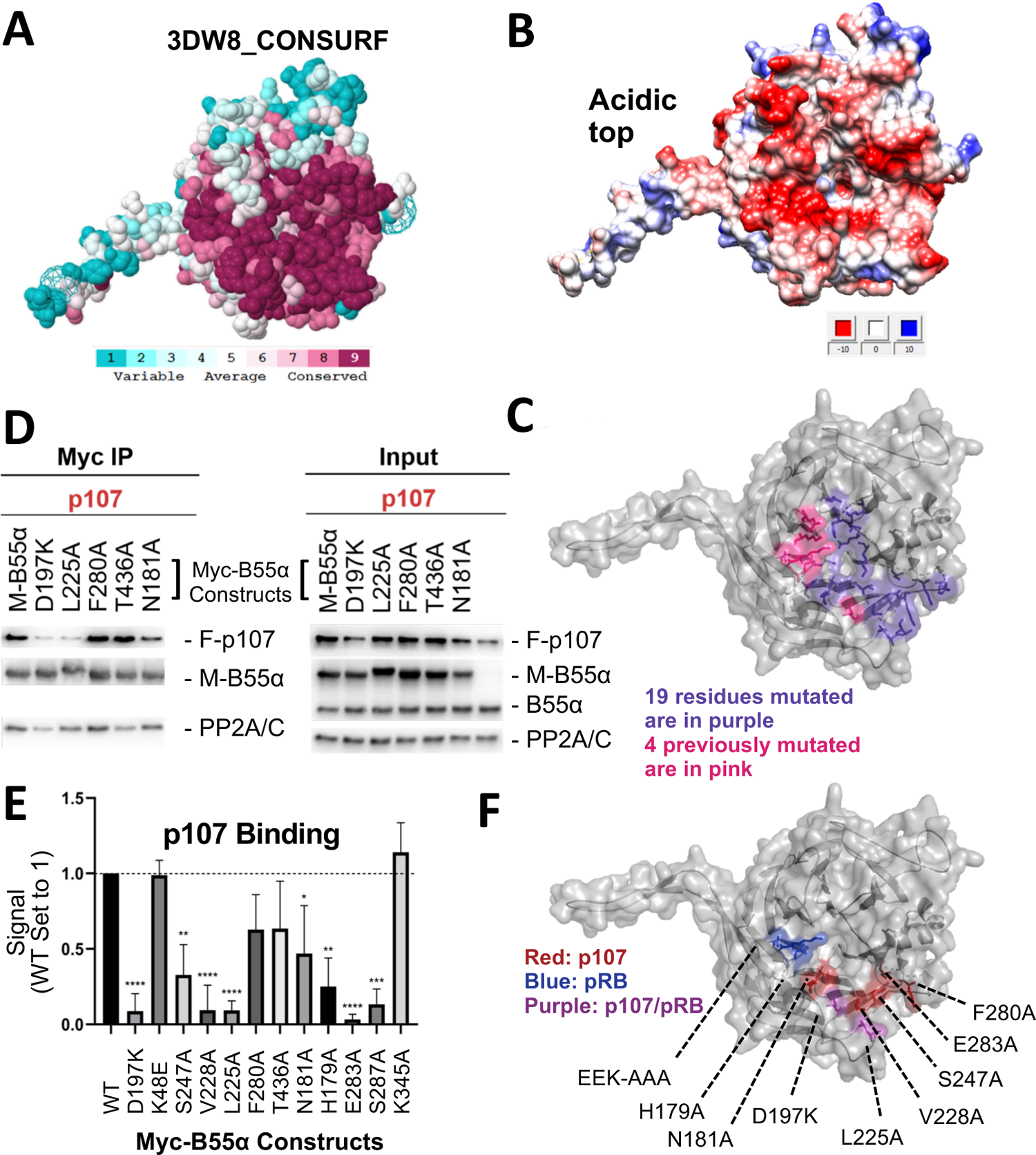
Mutation of highly-conserved residues on the β-propeller top of B55α have substrate specific effects on binding, supporting the notion that substrates contact different surfaces on B55α. (A) ConSurf depiction of PP2A/B55α mapping amino acid conservation (where amino acids are color-coded by conservation). (B) Electrostatic predictions mapped to the surface of the PP2A/B55α structure indicate an acidic top (red - acidic, blue - basic). (C) Nineteen single point mutations on the conserved top of the B55α β-propeller were generated. These are shown in purple. Four mutations generated and analyzed previously are shown in pink. (D) Representative immunoprecipitation experiment to test p107 binding requirements on Myc-B55α. Flag-p107 and wild-type and mutant Myc-B55α mutant constructs were co-transfected into HEK293T cells and used for IP’s with anti-Myc agarose conjugate. These assays were resolved via SDS-PAGE and proteins were detected using anti-Flag, anti-B55α, and anti-PP2A/C. (E) Mean values for cumulative immunoprecipitation assays for Flag-tagged p107 binding to Myc-B55α constructs are shown, with statistics indicated above. (F) The surface structure of B55α is depicted, with residues that appear important for p107 and pRB binding color-coded (in red and blue, respectively). The two Myc-B55α mutants that affect binding of both p107 and pRB (D197K and L225A) are colored purple.

For each substrate of interest, HEK293T cells were co-transfected with a FLAG-tagged substrate (p107, pRB or KSR1) and wildtype or mutant Myc-B55α constructs (Suppl. Fig 2B). Representative assays and the associated quantitation from the co-immunoprecipitation experiments are shown in Figs. 2D-E and Suppl. Fig 2C-D. B55α D197K and L225A variants showed the most profound impact on the recruitment of all three substrates when directly compared to wildtype B55α. Other B55α mutants showed varying effects on binding depending on the substrate, as illustrated by B55α V228A affecting only p107 binding while B55α H179A affected p107 and KSR1 but not pRB binding (Suppl. Fig 2C-D). To ensure that B55α variants did not induce a change in overall protein conformation, we assessed binding of all B55α mutants to PP2A/C. B55α D197K was the only variant that had a minor effect on PP2A/A and PP2A/C recruitment (Fig. 2D, left panel).

Taken together, while all substrates use the highly conserved top and groove of B55α for their interaction, our data demonstrate the contribution of distinct residues on the B55α surface for binding to different substrates (Fig. 2F). The two B55α residues required by the three substrates tested, D197K and L225A, are located on the B55α blade 4, which directly points towards the active site of PP2A/C. All other residues are an extension of this core binding interaction between the substrates and B55α.

### p107 S615 and S640 are PP2A/B55α dephosphorylation sites

PP2A/B55α is known to counteract phosphorylation by CDK kinases during mitosis. p107_M612-S687_ contains three SP sites (S615, S640 and S650, Fig. 1C) that can be phosphorylated in cells (Hornbeck *et al*, 2015). Using recombinant Cyclin A/CDK2 and γ-^32^P-ATP, we observed robust *in vitro* phosphorylation of p107. Our data suggested that at least two different sites are phosphorylated (Suppl. Fig. 3A). We confirmed these phosphorylation sites using antibodies raised against the phospho-CDK substrate motif “K/HpSP” (Suppl. Fig. 3B).

To further analyze PP2A/B55α-mediated dephosphorylation, we purified recombinant PP2A/B55α (Fig. 3A, *right panel*) (Zhao *et al*., 2019) and used this functional holoenzyme to analyze time-dependent dephosphorylation of p107_M612-S687_ (Fig. 3A, left panel), which is inhibited by 10 nM of the PP2A/C specific inhibitor okadaic acid. To determine which sites on p107 were dephosphorylated, we generated p107 mutants in which only a single “SP” site can be phosphorylated (i.e., S615A-S640A, S615A-S650A, and S640A-S650A), as well as a triple deletion control (S615A-S640A-S650A) (Fig. 3B). To validate phosphorylation/dephosphorylation and determine which sites were targeted by PP2A/B55α, we used mass spectrometry (MS). MS identified pS615 and pS640 in CDK2-phosphorylated p107 and a significant reduction of pS615 and pS640 phosphorylation upon incubation with PP2A/B55α (Fig. 3C). Interestingly, S650 phosphorylation was not identified, likely due to the close proximity of S650 to the RXL_658-660_ motif that mediates Cyclin A/CDK2 binding to p107.

**Figure 3.**
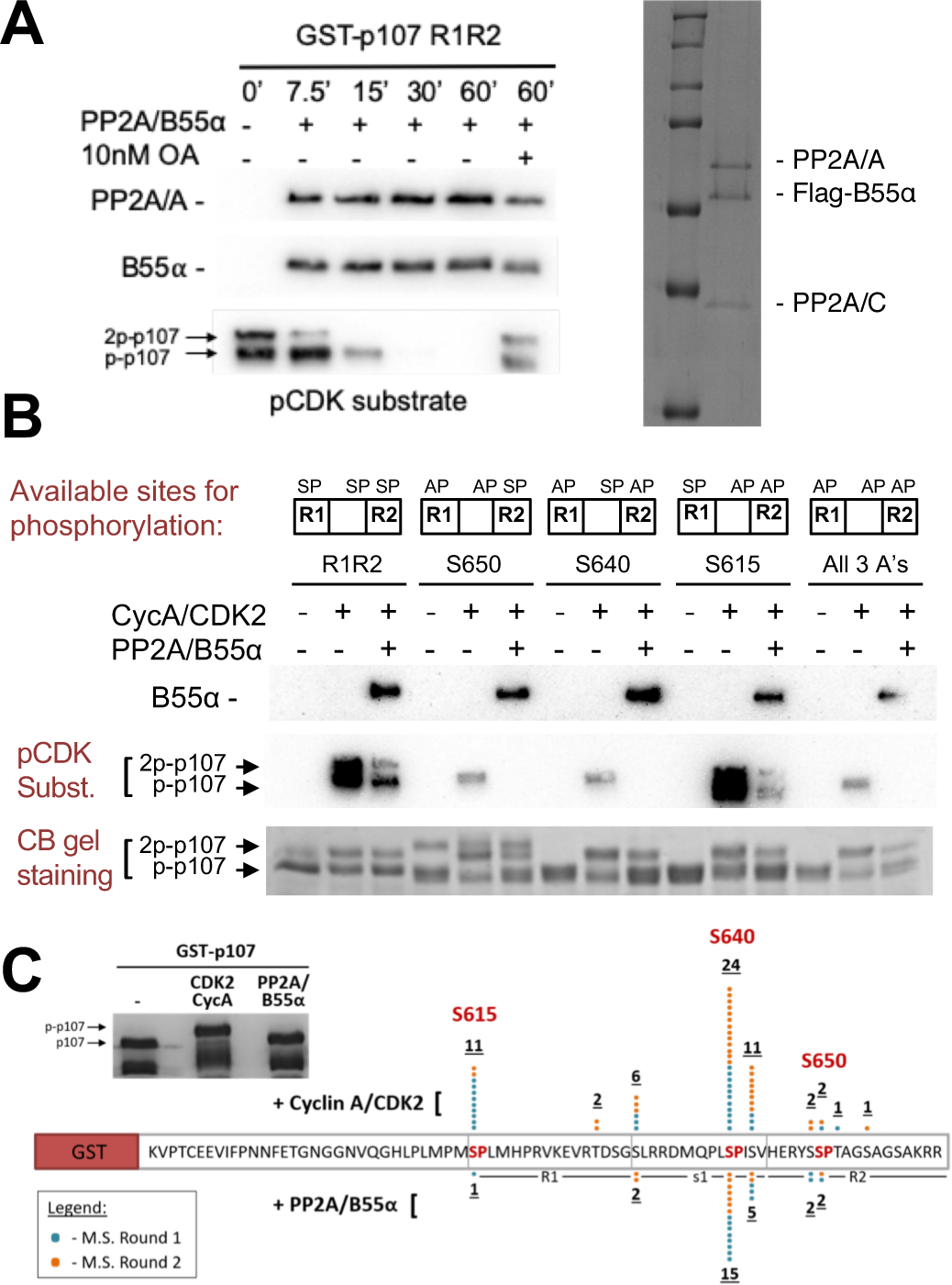
Combination of in vitro enzymatic assays and mass spectrometry identified S615 on R1 of p107 as the major site of PP2A/B55α-mediated dephosphorylation. (A) Dephosphorylation of GST-p107 R1R2 was performed using approximately 10 ng purified PP2A/B55α. The indicated time points were collected and samples were resolved via SDS-PAGE. Proteins were detected with anti-PP2A/A, anti-B55α and anti-pCDK substrate [(K/H)pSP]. Representative Coomassie Blue-stained gel depicting affinity purified PP2A/B55α holoenzymes is shown right. (B) In vitro phosphorylation and dephosphorylation of GST-p107 R1R2 with single SP sites available were performed using 0.25 μg recombinant Cyclin A/CDK2 and approximately 10 ng PP2A/B55α (each for one hour, respectively). Proteins were resolved via SDS-PAGE and detected by Coomassie gel staining and western blotting using anti-B55α and pCDK substrate antibodies. Note basal levels of pCDK substrate signal in all “+CycA/CDK2” lanes which indicates phosphorylation on CDK2 itself. (C) Representative Coomassie Blue-stained PAGE used for mass spectrometry, with schematic summarizing the findings from two independent rounds of analyses, is shown.

Next, we performed *in vitro* dephosphorylation experiments using p107 variants that are deficient in B55α binding due to elimination of positive charges in R1 or R1-R2. As expected, reduced dephosphorylation of p107 was detected for the binding-deficient p107 R1 [R621A/K623A] variant when compared to wt-p107, confirming our newly identified substrate binding residues (Fig. 4A). Dephosphorylation was further reduced for the p107-R1-R2 variant (R1 [R621A/K623A]-R2 [K657A/R659A]) (Fig. 4A), correlating well with the NMR-based binding affinities.

**Figure 4.**
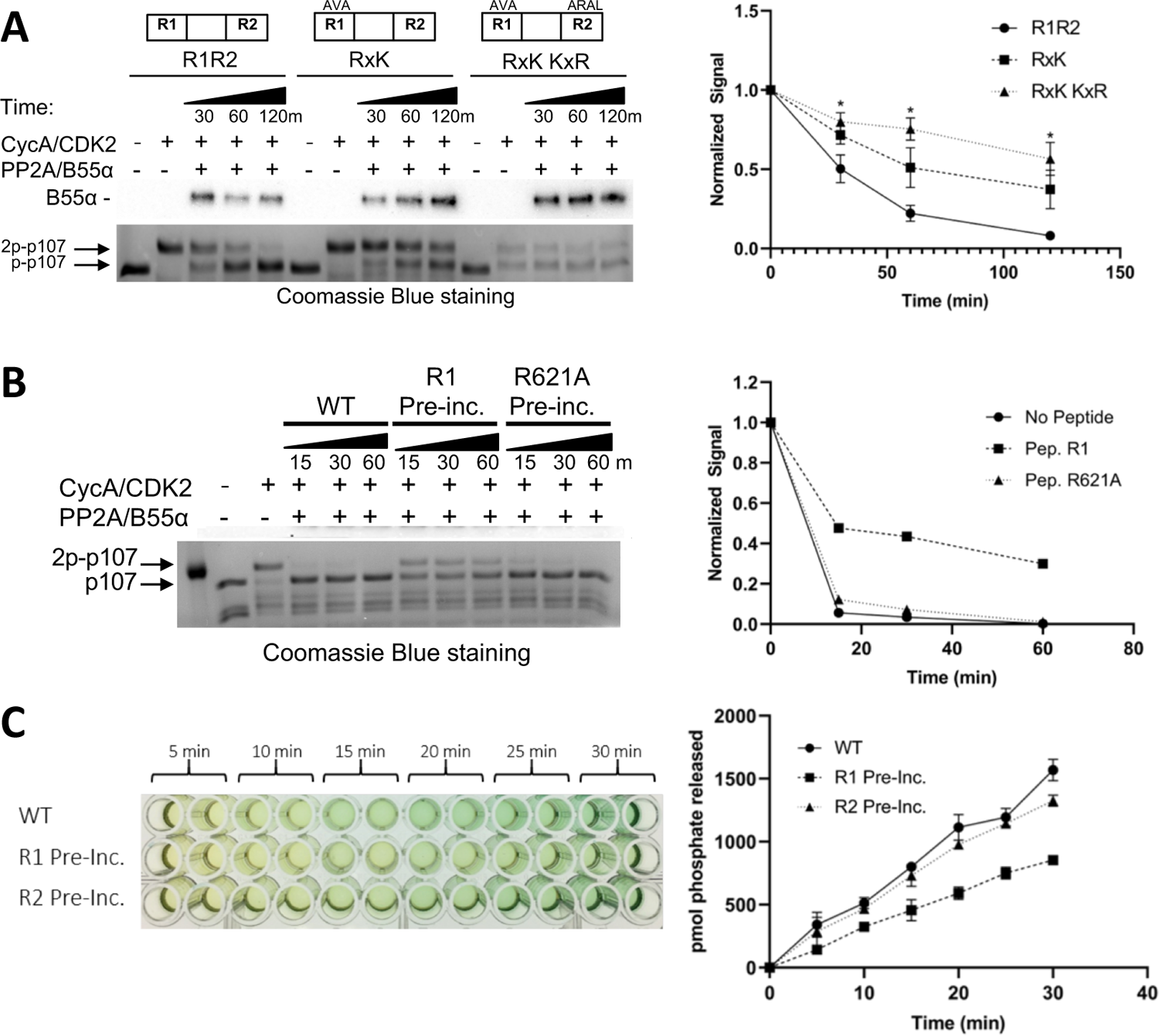
Residues critical for B55α/PP2A binding to p107 are critical for p107 spacer dephosphorylation. (A) Approximately 10 ng of purified PP2A/B55α was used to dephosphorylate wildtype GST-p107 R1R2 or mutant constructs (GST-p107 R621A K623A and GST-p107 RxK K657A R659A) in a time-course experiment. Proteins were resolved via SDS-PAGE and detected by Coomassie Blue staining or western blotting using anti-B55α antibodies. Quantifications of the “phospho”-p107 band for each construct tested were performed using ImageJ and plotted as a function of time, with statistics shown above. (B) Representative assay in which purified PP2A/B55α was preincubated with either wildtype R1 peptide or R621A mutant R1 peptide and then used in time-course dephosphorylation assays using GST-p107 R1R2 (native enzyme was used as a positive control for dephosphorylation). Proteins were resolved via SDS-PAGE and detected by Coomassie Blue. Quantification is shown right. (C) Purified PP2A/B55α was preincubated with either wildtype R1 or R2 peptide and then used in Malachite Green Phosphatase Assays with a p107-derived phosphopeptide (in which S615 is the available phosphosite). The indicated time points were collected and absorbance was read at 600 nm by microplate reader (quantification is shown right).

Lastly, we also assessed PP2A/B55α-mediated dephosphorylation of substrates using competitive peptides, specifically a peptide encompassing the R1 region of p107. As expected, dephosphorylation of p107 was significantly reduced when PP2A/B55α incubated with the R1 peptide was used for dephosphorylation studies (Fig. 4B). Repeating this experiment using a binding-deficient peptide (either R1 R621A or R2, respectively) showed dephosphorylation kinetics similar to that of the free PP2A/B55α, suggesting that the mutant peptide bound too weakly to compete for the substrate binding site on PP2A/B55α (Fig. 4B-C).

### p107 R1 residues that are critical for binding to PP2A/B55α and dephosphorylation

To understand which R1 residues are critical for the p107:B55α interaction we performed binding competition assays. In this assay we use increasing concentrations of R1 peptide to compete for B55α binding (Fig. 5A). This peptide was able to compete with a p107 R1R2 construct. As expected, using different peptides, such as R2 or small spacer peptides, did not show this effect. We performed this binding competition assay using endogenous PP2A/B55α from cellular extracts and observed no difference, highlighting that the source of PP2A/B55α did not matter (Suppl. Fig. 4A).

**Figure 5.**
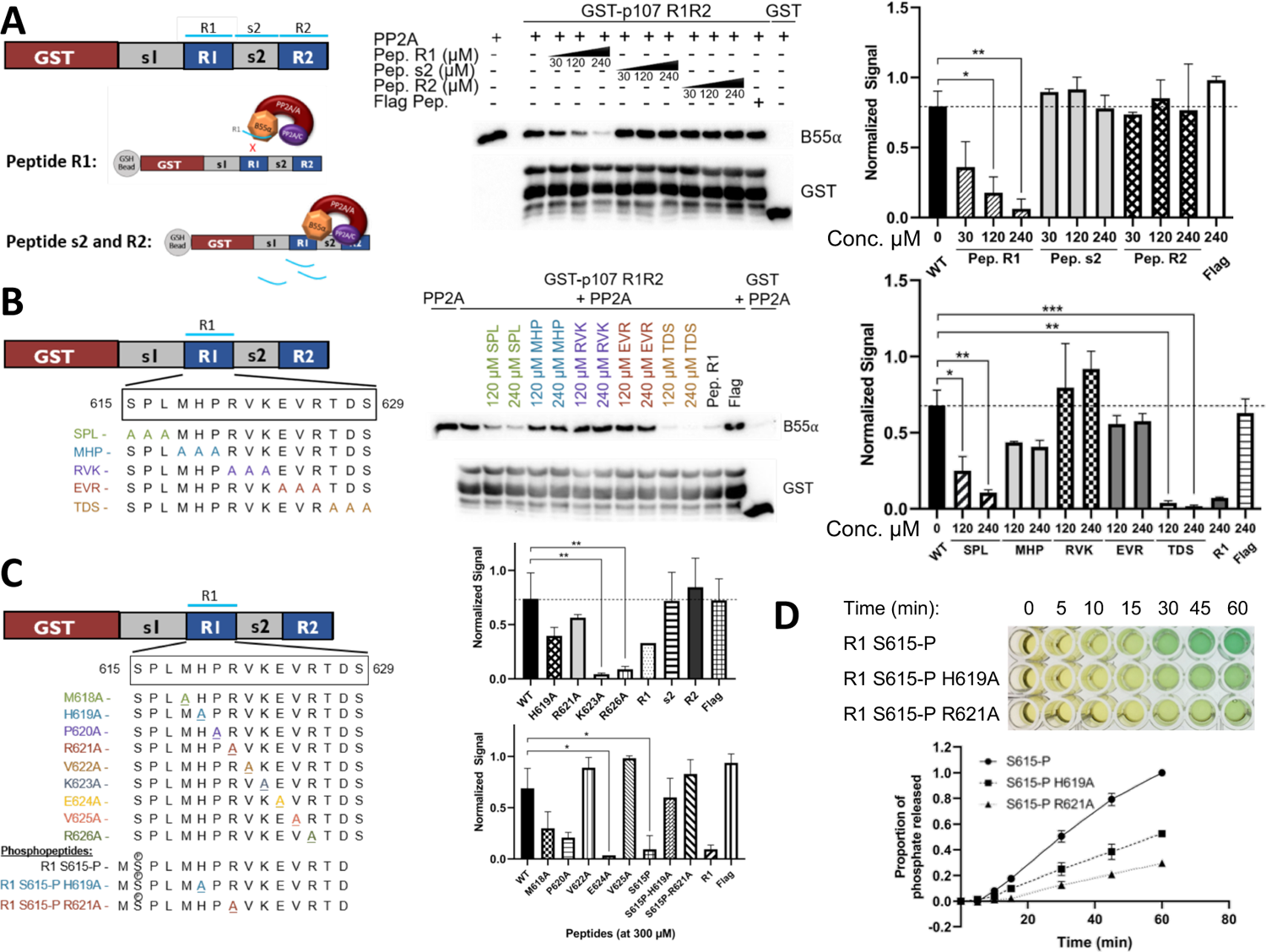
Identification of critical residues within the central 9-mer stretch of the “R1” region of p107 for binding to B55α/PP2A (SPxxHxRVxxV). (A) Purified PP2A/B55a was pre-incubated with synthetic p107 peptides and then used in pulldown assays with GST-p107 R1R2 constructs. Proteins were resolved via SDS-PAGE and detected via western blotting using anti-B55α and GST antibodies. Quantification of B55α pulled down relative to the wildtype pulldown was performed using ImageJ, with statistics shown above. (B-C) Pulldowns were performed using purified PP2A/B55α and GST-p107 R1R2 as above with pre-incubations using mutant p107-derived synthetic peptides (both scanning triple-mutants and point mutants) as well as wildtype phosphopeptides. Proteins were resolved via SDS-PAGE and detected via western blotting using anti-B55α and GST antibodies. Quantifications were performed as above. (D) Purified PP2A/B55α was used in time-course Malachite Green Phosphatase assays using wildtype or mutant p107-derived phosphopeptides as substrates of dephosphorylation. The indicated time points were collected and absorbance was read at 600 nm by microplate reader (quantification of duplicate assays is shown below).

Next, we generated a large battery of R1 peptides – all with single or multiple Ala substitution – and performed this binding competition assay. Our data clearly showed that three triple mutant peptides were unable to bind PP2A/B55α (Fig. 5B). These data showed that the main R1 interaction residues must be within p107 M618-R626. Next, we tested the single Ala substitutions peptides for p107 M618-R626 to identify all residues necessary for p107 binding to PP2A/B55α. These experiments showed that p107 residues H619, R621, V622, and V625 are critical for B55α binding, as R1 peptides with H619A, R621A, V622A, and V625A bound poorly to PP2A/B55α (Fig. 5C). Moreover, to confirm the requirement of each one of these residues for binding to B55α, we generated point mutants and tested them in binding assays. H619A, R621A, V622A, and V625A dramatically reduce binding to B55α holoenzymes (Suppl. Fig. 4C). These experiments defined a putative SLiM binding motif for p107 – HxRVxxV – with PP2A/B55α.

To assess the relationship between binding and dephosphorylation of p107 by PP2A/B55α directly, we measured dephosphorylation of wildtype (wt) pS615-R1 or mutant (mt) variant (H619A or R621A) phosphopeptides (all phosphopeptides acted as their unphosphorylated counterparts in competitive binding assays, Fig. 5C and Suppl. Fig. 4D). Fig. 5D shows that the mutant phosphopeptides were dephosphorylated by purified PP2A/B55α at a slower rate compared to the wildtype pS615-R1 phosphopeptide. These data provide us with direct evidence of delayed kinetics of PP2A/B55α-mediated dephosphorylation of a p107-derived substrate when binding is impeded.

Altogether, these binding and enzymatic assay data support a mechanism whereby the p107-based putative *HxRVxxV* SLiM docks at the mouth of the B55α groove (between blades 3 and 4 of the β-propeller) and the “SP” site within the active site of the PP2A/C catalytic subunit.

### B55〈 D197 plays a critical role in p107 recruitment

Next, we repeated our binding and dephosphorylation experiments in cells. Using co-transfection followed by immunoprecipitation assays, we show that GFP-p107-R1, but not mutant GFP-p107-R1 (H/AxR/AV/AxxV/A variant), immunoprecipitates both endogenous B55〈 and exogenous Myc-B55〈 together with PP2A/A (Fig. 6A). As expected, GFP-R1 did not immunoprecipitate B55〈-D197K (Fig. 6A). This confirms that R1 is sufficient to form a complex with the PP2A/B55〈 holoenzyme in cells and that this is dependent on B55〈 D197. The specificity of the GFP-p107-R1 fusion protein pull down was additionally validated by mass spectrometry, which resulted in the identification of B55α, B55δ, B55γ, B55β, PP2A/A and PP2A/C subunits, but no other PP2A or phosphatase subunits. Therefore, the p107 R1 domain contains a highly specific SLiM for B55〈 family members. Consistent with this result, GST-p107-R1 and GST-p107-R1R2 pulled wt B55α, but not the B55α D197 mutant, from 293T cell lysates (Suppl. Fig. 4B). Moreover, to determine if B55α modulates the phosphorylation state of CDK2 sites on p107 in cells, we transfected 293T cells with Flag-p107 alone or in combination with either wt B55α or mt B55α-D197K. Fig. 6B shows that wt B55α expression, but not mt B55α-D197K, leads to reduced phosphorylation of p107 on sites recognized by the CDK substrate antibody, which include p107 S615 (this is the only site recognized by this antibody in the p107 spacer, Fig. 3B). Altogether these data strongly suggest that B55α-mediated dephosphorylation of S615 in cells is dependent on specific residues C-terminal from the target dephosphorylation site that mediate contacts with B55α D197 and/or neighboring residues in B55α.

**Figure 6.**
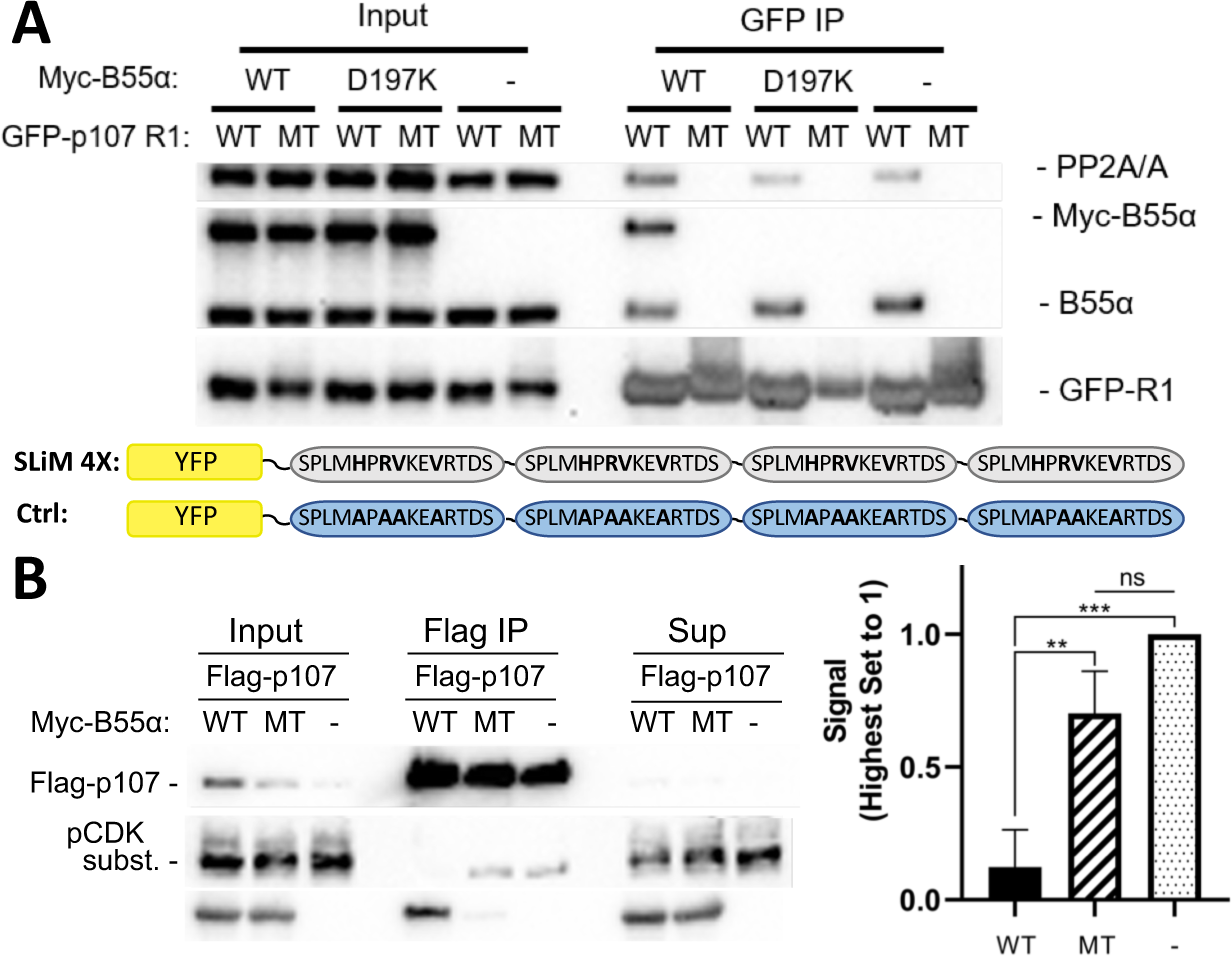
p107 R1 interaction with the B55α holoenzyme in cells depends on sites required for binding and dephosphorylation in vitro. (A) GFP-p107 R1 wild-type and mutant constructs were co-transfected with Myc-B55α wildtype and B55α-D197K mutant constructs into HEK293T cells and used for IP’s with anti-GFP agarose conjugate. Input lysates and IPs were resolved via SDS-PAGE and proteins were detected using anti-PP2A/A, anti-B55α, and anti-GFP. Schematic of WT and MT GFP-p107 R1 constructs is shown below. (B) Flag-p107 was co-transfected with Myc-B55α wildtype and B55α-D197K mutant constructs into HEK293T cells and used for IP’s with anti-Flag agarose conjugate. IPs and input/supernatant lysates were resolved via SDS-PAGE and proteins were detected using anti-Flag and anti-pCDK substrate antibodies. (Right) Quantification of pCDK substrate signal (relative to Flag) is shown, with statistics shown above.

### A model for p107 recruitment by PP2A/B55α

Based on all the established data, we created a model to describe the recruitment of p107 by PP2A/B55. This model is based on the following premises: 1) Using HxRVxxV as a ruler we assumed that either H619_p107_ or R621_p107_ interacts with D197_B55α_; and 2) as we confirmed that pS615_p107_ is dephosphorylated by PP2A/B55α, pS615_p107_ must be placed into the PP2A active site. In order to place pS615_p107_ it became more likely that R621_p107_ interacts with D197_B55α_ for simple distance reasons. This meant that pSPMLHPR will need to connect the PP2A active site and D197_B55α_. To identify possible existing confirmations of such a peptide we searched the PDB database for peptide fragments from existing proteins structure using the sequence EPxxxPR (pS was exchanged to the E mimic to identify more peptides). Next, we examined the distribution of distances between the oxygen atoms of the OE_1_/OE_2_ atoms of the E side chain and the NH_1_/NH_2_ atoms of the R side chain, and compared these to the distance between E in the PP2A active site and R621_p107_ bound to D197_B55α_, which is 27.5 Å (Fig. 7A-B, Suppl. Fig 5A). The 520 identified peptides with the sequences xEPxxxPRx have distances between 15-25 Å (Suppl. Fig. 5A), indicating that p107 pSPMLHPR must form a highly extended structure when bound to PP2A/B55. Lastly, we used an extended EPxxxPR peptide, placed it into the PP2A/B55α complex structure, mutated the EPxxxPR sequence to the p107 sequence (MpSPMLHPRV) and refined these structures using FlexPepDock (London *et al*, 2011; Raveh *et al*, 2010; Raveh *et al*, 2011).

**Figure 7.**
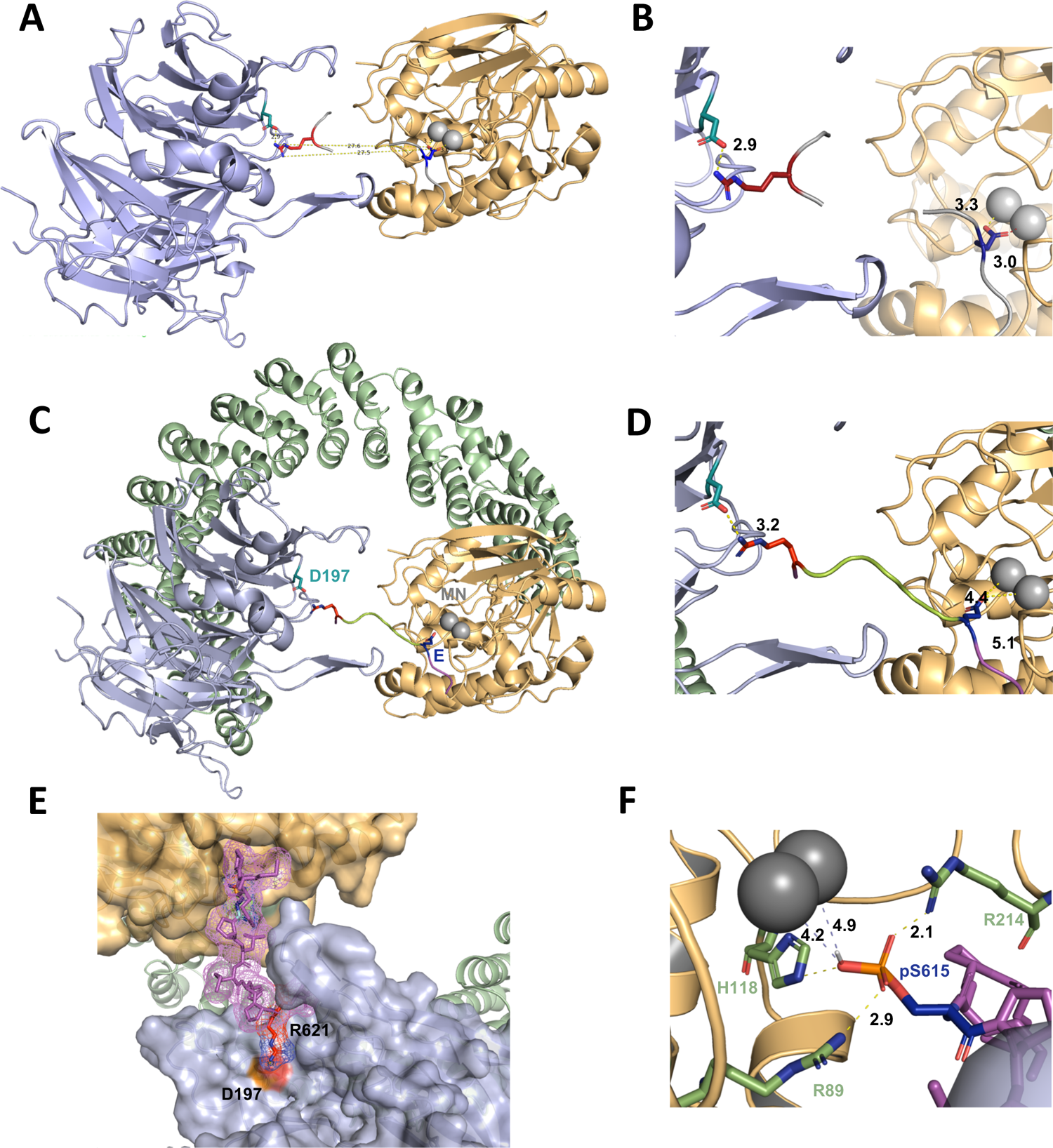
A computational model of the p107 phosphopeptide (613-622) binding B55α and the active site of PP2A/C is consistent with p107 spacer contacts to B55α as determined by NMR spectroscopy. (A) The “two-fragment” model depicting B55α and PP2A/C with two modeled peptide fragments. Distances between the OE1/OE2 atoms of Glu and the NH1/NH2 atoms of Arg are shown center. (B) A closer view of “two-fragment” model highlighting the distances between Arg and D197 of B55α and between the Glu residue and the Mn2+ ions within PP2A/C active site. (C-D) Peptide model depicting the best alignments with the two reference fragments and the best distances to the Mn2+ ions and D197. A closer view highlighting the distances between the NH2 of Arg residue and D197 of B55α, and the Glu residue with the Mn2+ ions, is shown in (C). (E) Close-up of model in (C-D) showing the peptide side chains and contacts to B55α surface. (F) Close-up of pSer-621 (substituted for the Glu residue) and residues critical for phosphate coordination (R630, H559, R655). Distances between residues where H-bonding is predicted to occur are shown.

In the best PP2A/B55α/p107 model, the NH_2_ side chain of R621_p107_ is 3.2 Å apart from D197_B55α_, while the OE_1_/OE_2_ of E615 (pS615 mimic) is 4.4 and 5.1 Å apart from the two Mn^2+^ ions at the PP2A active site, respectively (Fig. 7C-E). These distances are slightly longer than expected for this type of interactions, but replacing E615 with pS615 show consistency with active site binding of PPPs (Fig. 7F) (Goldberg *et al*, 1995). Taken together, our model shows that p107 pSPLMHPRV can bind in a highly extended fashion to the PP2A/B55α holoenzyme, in full agreement with our experimental data.

### A derived p107-pSPxxHxRVxxV-SLiM is conserved in other substrates and functional validated in TAU

Peptides known to abrogate TAU and MAP2 binding to B55α have been reported (Sontag *et al*, 2012). We noticed that TAU and MAP2 and the conserved p107 family member, p130, contain residues that align with our defined p107 SLiM, generating a consensus sequence, **p**[**ST**]-**P-**x(5-10)-[**RK**]-**V**-x-x-[**VI**]-**R** (Fig. 8A) and other cellular IDPs, where the phosphosite is 5-10 residues amino terminal from the conserved residues in the binding motif. To functionally validate the conservation of the p107 SLiM we selected TAU, a well know B55α/PP2A substrate, which has been shown to require acidic residues on the B55α surface for binding, but any other residues that form a defined SLiM are unknown. Therefore, we generated a phospho-TAU peptide encompassing the putative conserved SLiM and a variant peptide with the conserved residues mutated to Ala. Fig. 8B shows that disruption of critical residues within the newly defined B55α substrate SLiM completely block TAU dephosphorylation.

**Figure 8.**
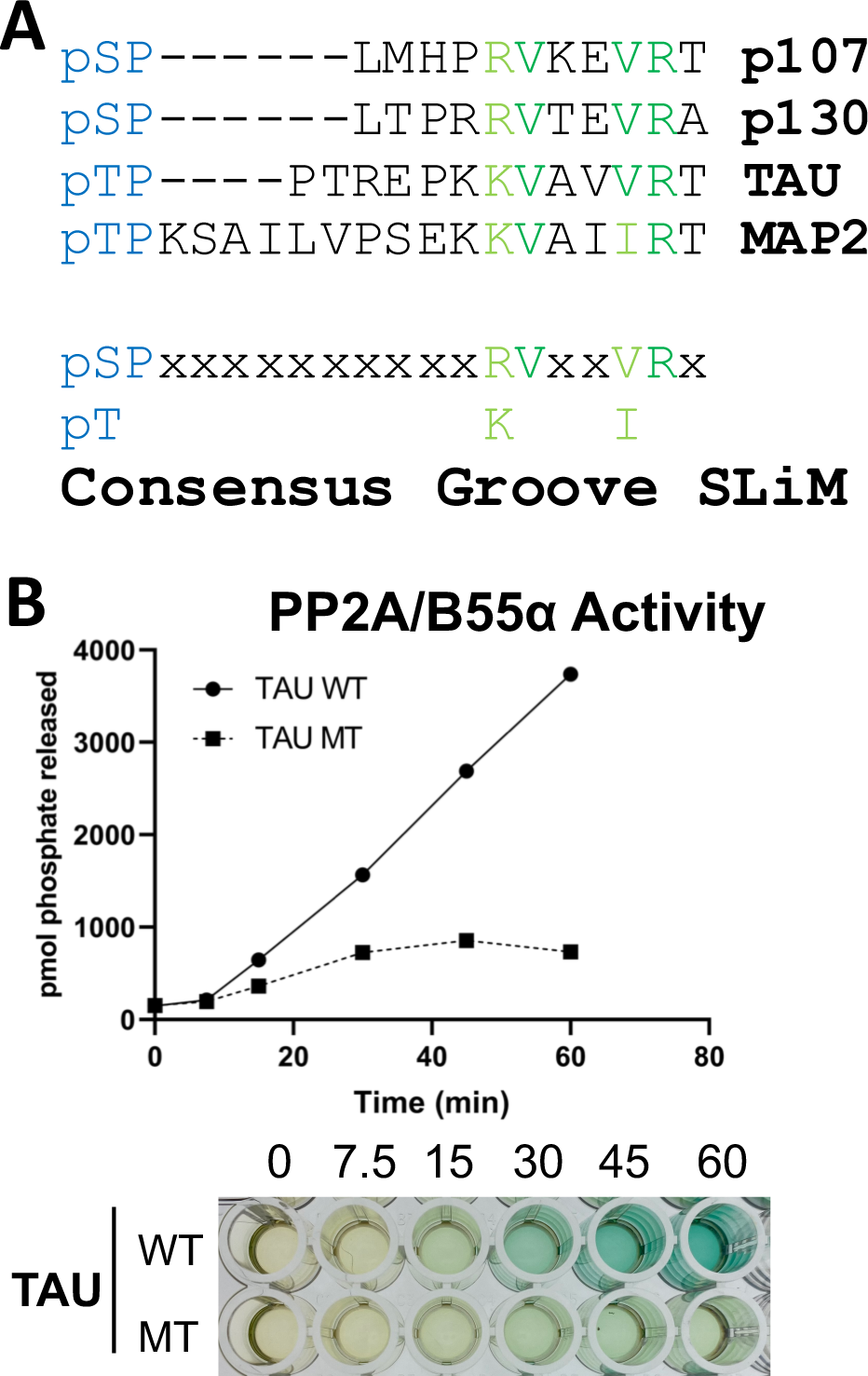
A derived p107-pSPxxHxRVxxV-SLiM is conserved in other substrates and functional validated in TAU. (A) Schematic of our proposed hypothetical consensus groove SLiM, p[ST]-P-x(5-10)-[RK]-V-x-x-[VI]-R, for TAU, MAP2, and the conserved p107 family member, p130, each of which contain residues that align with our defined p107 SLiM. (B) Time-course Malachite Green Phosphatase assay using a wild-type phospho-TAU peptide encompassing the putative conserved SLiM and a variant peptide with the conserved residues mutated to Ala. (Above) Quantification of phosphatase assay shown below.

## Discussion

PP2A is a key Ser/Thr protein phosphatase, which requires its B regulatory subunits to specifically recruit substrates. PP2A B-subunits are proteins that adopt specific folds and thus allow for specific substrate recruitment. Much work over the last few years has highlighted how the B56 regulatory subunit recruits its substrates. A central, conserved groove region in B56 provides the necessary and sufficient interaction site for the substrate recognition sequence (SLiM site) – LxxIxE (Hertz *et al*., 2016; Wang *et al*., 2016). Much less is known for B55, despite the fact that it is the most abundant B subunit and targets a myriad of key cellular substrates. Previous reports highlighted the possibility that B55〈 functions differently with the potential presence of distinct substrate recruitment sites. Specifically, sites of the acidic surface of B55α that were required for Tau dephosphorylation (Xu *et al*., 2008) appeared dispensable for p107 binding (Jayadeva *et al*., 2010). Here we used a broad range of approaches – from molecular to biochemistry to cellular – to understand how PP2A/B55α recruits specific substrates, using p107 as a model substrate.

We show that p107 S615 (CDK kinase phosphorylation site) is a specific B55α/PP2A holoenzyme dephosphorylation site. p107 residues H619, R621, V622 and V625 within the p**S**Pxxx**H**x**RV**xx**V** motif are critical for binding to B55α. Next, we showed that these residues bind to a B55α surface groove that is defined by residues D197 and L225. Importantly, our results show that this B55α surface is also important for recruitment of pRB and KSR, further suggesting that these sites are likely the key substrate recruitment sites for a variety of substrates. However, pRB and KSR differ in the requirement of other conserved residues present within the B55α groove, strongly suggesting that while substrate specificity is determined by the groove, it may use different residues to accommodate a variety of substrates. With this regard, we have found that peptides known to abrogate Tau and MAP2 binding to B55α (Sontag *et al*., 2012) contain residues that align with our proposed SLiM consensus sequence, **p**[**ST**]-**P-**x(5-10)-[**RK**]-**V**-x-x-[**VI**]-**R,** that is also found in p130/RBL2 (Fig. 8A) and other cellular IDPs, where the phosphosite is 5-11 residues amino terminal from the conserved residues in the binding motif. Strikingly, purified B55α/PP2A holoenzymes dephosphorylate the wildtype TAU peptide, but fail to dephosphorylate a mutant variant that lacks the conserved residues. It appears that the number of residues between the conserved residues that contact the groove and the phosphorylation site can vary but it is small (5 residues in p107 vs. 7 residues in TAU and 11 in MAP2), which suggests a role for the SLiM in phosphosite presentation. In contrast, recent work on the B56 SLiM suggests that B56 SLiM-containing proteins are direct substrates as well as scaffolds to facilitate the recruitment of proteins to B56/PP2A for dephosphorylation (Kruse *et al*, 2020). While there is some correlation between the distance of the B56 SLiM and the site of dephosphorylation and the rate of dephosphorylation, many B56-dependent phosphorylation sites are located on non-SLIM containing proteins indicating B regulatory subunit specific dephosphorylation mechanisms.

Moreover, we also identified a second region of p107, which we termed R2, that contributes to B55α binding. This R2 region includes a higher density of positively charged residues that enhance binding most likely via charged:charged interactions. Indeed, this has also been recently shown for protein phosphatase 2B (PP2B; PP3; Calcineurin) and the B56 regulatory subunit of PP2A, where these dynamic charged:charged interactions were shown to play a role in substrate specificity (Hendus-Altenburger *et al*, 2019; Wang *et al*., 2020). Thus, such interactions, which have been recently recognized to be critical for the interaction of IDPs with their target proteins and allow for increased entropy, are likely important for substrate recruitment and specificity by different phosphatases.

Lastly, we leveraged our mutagenesis data to understand the p107 R1 binding site on B55α by generating a model how p107 R1 engages simultaneously with B55α and the PP2Ac active site. Our computational model based on structural data of peptides with the conserved key residues on p107 needed for B55α binding shows the feasibility of simultaneous binding of both the active site of PP2A/C and D197 on the B55α groove. However, the model peptide is significantly extended, a conformation that would not be favored for catalysis without additional bending of the scaffold subunit. Analysis of the flexibility of the PP2A/A scaffold upon binding to the catalytic subunit, and B subunits of the four distinct holoenzymes, confirms that subunit binding has a major effect on PP2A/A bending (Suppl. Fig. 6). The dependency of the conformational flexibility of the scaffold subunit on the nature of the B subunit bound strongly suggests that substrate binding should result in additional relative movement of PP2A/A HEAT-repeats to promote catalysis.

Taken together, our detailed molecular and cellular study highlights how PP2A/B55α recruits substrates and how these substrates engage with the PP2A active site, and uncovers the top groove of B55α as a central hub for substrate discrimination. In addition, this work will also facilitate identification and validation of new substrates based on the presence of variant SLiMs, a key step towards understanding B55α/PP2A role in multiple cellular processes.

## Materials and Methods

### Cell culture and cell lines

All cell lines were obtained from ATCC and cultured in DMEM supplemented with 10% FBS and 0.1% Penicillin-Streptomycin as described previously (Jayadeva *et al*., 2010) and tested for mycoplasma annually. For stable expression of Flag-B55α, HEK293T cells were transfected with pCPP-puro-Flag-B55α followed by puromycin selection for clone generation. Transient expression of Myc- and Flag-containing constructs was achieved using calcium phosphate transfection. Briefly, 5 µg plasmid DNA added dropwise to 2X HEPES-buffered saline (HBS) solution (280 mM NaCl, 50 mM HEPES, 1.5 mM Na2HPO4, pH 7.05) with bubbling, followed by addition to cells treated with 25 mM chloroquine after a 30-minute incubation period.

### GST Pulldown Assays

GST-tagged constructs of interest were expressed in *E. coli* bacteria and purified for use in pull-down assays. Briefly, 100 mL cultures of *E. coli* were treated with 0.25 mM IPTG for 2 hours to induce expression of GST-fusion proteins. Cells were then harvested by centrifugation and resuspended in NETN lysis buffer (20 mM Tris pH 8, 100 mM NaCl, 1 mM EDTA, 0.5% NP-40, 1 mM PMSF, 10 µg/mL Leupeptin) prior to sonication at 30% amplitude for 10 cycles. Supernatants were collected and incubated with glutathione beads for purification, followed by NETN buffer washes for sample clean-up. For pull-down assays, purified GST-p107 spacer and mutant constructs were incubated with HEK293T lysates for 3 hours or overnight at 4 °C, followed by washes (5X) with complete DIP lysis buffer (50mM HEPES pH 7.2, 150 mM NaCl, 1 mM EDTA, 2.5 mM EGTA, 10% glycerol, 0.1% Tween-20, 1 µg/mL aprotinin, 1 µg/mL leupeptin, 1 µg/mL Pepstatin A, 1 mM DTT, 0.5 mM PMSF) and elution with 2X LSB. Samples were resolved by SDS-PAGE and probed using antibodies against proteins of interest.

### Peptide Competition Assays

Synthetic peptides derived from the amino acid sequence of p107 were generated and used in competition assays with GST-tagged p107 R1R2 constructs (Biomatik). Briefly, purified PP2A/B55α holoenzymes were pre-incubated with 30-300 µM synthetic p107 peptides for 30 minutes at 37 °C to facilitate interaction. These pre-incubated PP2A/B55α complexes were then incubated with GST-tagged p107 R1R2 constructs for 3 hours or overnight at 4 °C, followed by washes (5X) with NETN lysis buffer and elution with 2X LSB. Proteins were resolved via SDS-PAGE and detected via western blotting using anti-B55α and GST antibodies. Densitometric quantitation was performed using ImageJ software.

### Immunoprecipitations

Whole cell extracts (200-400 µg) were incubated with Myc- or Flag-conjugated beads (Sigma) for 3 hours or overnight at 4 °C. Input samples were collected prior to antibody-conjugated bead incubation, and supernatants were taken post-incubation following sample centrifugation. Beads were then washed (5X) with complete DIP lysis buffer and proteins were eluted in 2X LSB. Proteins were resolved by SDS-PAGE and probed using antibodies against proteins of interest.

### PP2A/B55α Purification

Stably-expressing Flag-B55α HEK293T cells were expanded into six 15-cm tissue culture plates until they reached confluency. Cells were then harvested, washed 3X in cold 1X PBS, and then lysed in complete DIP lysis buffer for one hour. Lysates were then incubated with Flag-conjugated beads for 5 hours or overnight at 4 °C. After washing the beads 8X with complete DIP lysis buffer, elution buffer containing 200 µg/mL DYKDDDDK peptide (Genscript) was added to samples and incubated with shaking 2X for 30 minutes each (eluates were collected after each incubation). Eluates containing purified PP2A/B55α complexes were combined with 25% glycerol and 1 mM DTT for −80 °C storage.

### *In vitro* Kinase and Phosphatase Assays

GST-tagged p107 constructs loaded on beads were phosphorylated with recombinant Cyclin A/CDK2 (Invitrogen) in 200 nM ATP KAS buffer (5 mM HEPES pH 7.2, 1 mM MgCl2, 0.5 mM MnCl2, 0.1 mM DTT). We incubated samples with shaking at 37 °C for 2 hrs unless indicated. Reactions were stopped by adding 2X LSB and heated at 65 °C when used for PAGE or immunoblots. *In vitro* phosphorylated substrates for phosphatase assays, were washed 3X in complete DIP buffer, followed by addition of indicated concentrations of purified PP2A/B55α. Reactions were incubated with shaking at 37 °C for times indicated, followed by addition of 2X LSB and boiling at 65 °C for western blotting.

### Malachite Green Phosphatase Assay

The phosphatase activity of purified PP2A/B55α complexes was assessed using a commercial Ser/Thr phosphatase assay kit (EMD Millipore). Briefly, phosphatase-containing samples were incubated with synthetic p107-derived phosphopeptides for various time points at RT (up to 30 minutes). Malachite green reagent with 0.01% Tween-20 was then added to quench the reaction. Absorbance was determined at 600 nm on a microplate reader.

### Molecular Modeling of a p107 peptide presented by the groove on B55α to the active site of PP2A/C

To dock R621 to B55α, we retrieved peptide structures with “xHPRVx” from the PDB, then calculated dihedral angles (ϕ, φ and ω) of each residue of HPRV motif. The distance of each pair of peptide structures is the maximum distance of dihedral angles of residue pairs as previously described (North *et al*, 2011). The peptide structures were clustered by DBSCAN function in the R project (https://cran.r-project.org/web/packages/dbscan/index.html). We used the structure with best resolution (1G8M and chain A, resolution = 1.75 Å) in the largest cluster as our reference peptide. The peptide structure was moved to D197 of B55α in PDB: 3DW8 by manual rotations and translations in PyMOL, such that the NH1/NH2 atoms of R621 were approximately 3Å from OD1/OD2 atoms of D197. This model was subsequently refined with FlexPepDock web server (http://flexpepdock.furmanlab.cs.huji.ac.il/) (Fig. 7A-B).

To identify the likely phosphate position in the PP2A active site, we retrieved 68 structures from the PDB containing a phosphate group (PO_4_) and a pair of Mn^2+^ ions from a search on the Mn^2+^ ion on our web site Protein Common Interface Database (ProtCID), http://dunbrack2.fccc.edu/ProtCiD/IPdbfam/PfamLigands.aspx?Ligand=MN) (Xu & Dunbrack, 2020). These 68 structures contain 47 distinct UniProt proteins including four human phosphatase proteins (PP1A_HUMAN: PDB: 4MOV, 4MOY and 4MP0, PP1G_HUMAN: PDB: 4UT2 and 4UT3, PPP5_HUMAN: PDB: 1S95 and PR15A_HUMAN: PDB: 4XPN). For each Mn^2+^ ion, we calculated the distances from the four oxygen atom of the phosphate, and saved the shortest distance. The average distance from PO_4_ to Mn ions is 2.38 Å and the standard deviation is 0.59 Å. We also examined a structure of PP5 (PDB: 5HPE) that contains a substrate peptide (expressed as a tag separated by a linker from the C-terminus of PP5) with a Glu side chain as phosphomimetic. After aligning the proteins homologous to PP2A with bound phosphate, the structure of 5HPE with a Glu residue, and PP2A in PDB: 3DW8, it is clear that the oxygen atoms of the phosphates and the Glu are in very similar positions relative to the Mn^2+^ ions (Suppl. Fig. 5D). We defined a structure of PP2A+B55α with the Glu phosphomimetic from PDB: 5HPE and the HPRV peptide as “the two fragment model.”

To build p107 peptide (MPMEPLMHPRV), we searched the PDB and found 520 peptide structures that contain the sequence “EPxxxPR” which would connect phosphate (with E as a phosphomimetic of pSer) and Arg in the two binding sites. We calculated the distances between the OE1/OE2 of Glu and NH1/NH2 of Arg for these 520 structures to find peptides long enough to connect Glu and Arg residues. We superposed 217 peptide structures with a distance ≥ 20 Å onto the “two-fragment model” by minimizing the distances of the OE1/OE2 and NH1/NH2 atoms of the peptide with those of the two-fragment model by *pair_fit* function in PyMOL (Suppl. Fig. 5B), then calculated the sum of the displacements of the OE and NH atoms for each peptide from the equivalent atoms in the two-fragment model (Suppl. Fig 5C). The top 20 structures with minimum distances to the OE1/OE2 atoms of Glu and NH1/NH2 of Arg residues within the reference peptide fragments were selected and mutated into “EPLMHPR”, and added “MPM” residues and “V” residue to build the p107 peptide “MPMEPLMHPRV” in PyMOL (Suppl. Fig 5E). Each PP2A-B55α-p107 structure was refined with the FlexPepDock server and the top 10 models with the best Rosetta energy scores were saved (Alford *et al*, 2017). The model in which the OE1/OE2 atoms of Glu and NH1/NH2 atoms of Arg of p107 peptide has the minimum distances to the Mn^2+^ ions and D197 was selected. To assemble our PP2A-p107 complex model, we then superposed 3DW8 onto this model to add the scaffold PP2A/A subunit (Fig. 7C-D).

### NMR Spectrometry

#### Plasmid Construction for NMR studies

B55α_M1-N447_ gene was subcloned into a modified pCDNA3.4 vector (pcDNA3.4-K-GFP-RP1B). The pcDNA3.4-K-GFP-RP1B vector contains a Kozak consensus sequence, an N-terminal His6-green fluorescent protein-tag, a TEV cleavage site and a multiple cloning site following the pcDNA3.4 human cytomegalovirus (CMV) promoter.

This plasmid was amplified and purified using the NucleoBond PC 500 Plasmid Maxiprep Kit (MACHEREY-NAGEL). P107_M612-S687_ was subcloned into an MBP-fusion vector (pTHMT). The PTHMT vector contains an N-terminal His6-Maltose Binding Protein (MBP)-tag, a TEV cleavage site and a multiple cloning site following the T7 promoter.

#### Protein Expression and purification for NMR studies

B55α_1-447_ was expressed in Expi293F cells (ThermoFisher) at a ratio of 1.0 µg DNA per mL of final transfection culture volume. Transfections were performed using 125 ml medium (Gibco Expi293 Expression Medium) in 250 ml baffled flasks (Corning) according to the manufacturer’s protocol in a humidified incubator at 37 °C and 8.0% CO_2_ under shaking (125 rpm). On the day of transfection, the cell density was between 3-5 × 10^6^ cells ml^−1^. Prior to transfection, the Expi293F cells were seeded at 2.6 × 10^6^ cells ml^−1^ in 85% of the final transfection volume. B55α DNA was mixed with Opti-MEM Reduced Serum Medium (ThermoFisher); in a separate tube, polyethylenimine (PEI) reagent was mixed with Opti-MEM medium. The DNA and PEI mixtures were combined and incubated for an additional 10 min. The final transfection mixture was then added to the cells. 2 mM (final concentration) valproic acid was added to the cells 18 h post-transfection. After an additional 24–28 h, the cells were harvested and the pellet was stored at −80°C. P107_612-687_ was expressed in *E. coli* BL21 DES cells (Agilent). Cells were grown in Luria Broth in the presence of the selective antibiotic at 37 °C to an OD_600_ of ∼0.8, and expression was induced by addition of 1 mM isopropyl β-D-thiogalactoside (IPTG). Induction proceeded for 5 h at 37 °C prior to harvesting by centrifugation at 6,000 xg for 15 min (ThermoFisher). Cell pellets were stored at −80 °C until purification.

p107 cell pellets were resuspended in ice-cold lysis buffer (50 mM Tris pH 8.0, 0.5 M NaCl, 5 mM imidazole, 0.1% Triton X-100 containing a EDTA-free protease inhibitor tablet [Sigma]), lysed by high-pressure cell homogenization (Avestin C3 Emulsiflex) and centrifuged (35,000 xg, 40 min, 4 °C). The supernatant was loaded onto a HisTrap HP column (GE Healthcare) pre-equilibrated with Buffer A (50 mM Tris pH 8.0, 500 mM NaCl and 5 mM imidazole) and was eluted using a linear gradient of Buffer B (50 mM Tris pH 8.0, 500 mM NaCl, 500 mM imidazole). Fractions containing the protein were pooled and dialyzed overnight at 4 °C (50 mM Tris pH 7.5, 500 mM NaCl) with TEV protease to cleave the His_6_-MBP-tag. The cleaved p107 was heated at 80 °C for 10 min and centrifuged (29,000 xg, 20 min, 4 °C). The supernatant was concentrated to 3-4 ml and heated at 80°C for 10 min and centrifuged prior to SEC purification (SEC buffer: 20 mM sodium phosphate pH 6.5, 50 mM NaCl or 250 mM NaCl, 0.5 mM TCEP; column: Superdex 75 16/60, GE Helathcare) B55α cell pellets were resuspended in ice-cold lysis buffer containing EDTA-free protease inhibitor tablet (Sigma) and rocked at 4 °C for 20 min before centrifuged (42,000 xg, 50 min, 4 °C). The supernatant was mixed with GFP-nanobody coupled agarose beads (AminoLink Plus Immobilization Kit, Thermo Fisher Scientific) and rocked at 4 °C for 2 h, and centrifuged (1,000 xg, 5 min, 4 °C). The resin was washed twice with buffer (20 mM Tris, pH 7.5, 100 mM sodium chloride 0.5 mM TCEP) by centrifugation. The His_6_-GFP-tagged B55α loaded resin was then suspended in 20 ml of the wash buffer and incubated with TEV protease overnight at 4°C. The cleaved product was centrifuged, and the supernatant was collected. The protein was further purified using anion exchange chromatography (Mono Q 5/50 GL, GE healthcare) pre-equilibrated with ion exchange buffer A (20 mM Tris, pH 7.5, 100 mM sodium chloride 0.5 mM TCEP) and eluted using a linear gradient of buffer B (20 mM Tris, pH 7.5, 1 M sodium chloride 0.5 mM TCEP). Fractions containing B55α were identified using SDS-PAGE, pooled, and concentrated to designated concentration for experiments or stored at −80°C.

#### NMR Spectrometry Data Collection and Processing

NMR data were recorded at 283 K on a Bruker Neo 600 MHz or 800 MHz (^1^H Larmor frequency) NMR spectrometer equipped with a HCN TCI active z-gradient cryoprobe. NMR measurements of p107_612-687_ were recorded using either ^15^N- or ^15^N,^13^C-labeled protein at a final concentration of 6 or 200 μM in NMR buffer (20 mM sodium phosphate pH 6.8, 250 or 50 mM NaCl, 0.5 mM TCEP) and 90% H_2_O/10% D_2_O. Unlabeled B55α and ^15^N-labeled p107 was mixed at 1:1 ratio to form the complex. The sequence-specific backbone assignments of p107_612-687_ were achieved using 3D triple resonance experiments including 2D [^1^H,^15^N] HSQC, 3D HNCA, 3D HN(CO)CA, 3D HN(CO)CACB, 3D HNCACB, 3D HNCO, and 3D HN(CA)CO. All NMR data were processed using Topspin 4.0.5 and analyzed using Cara. All NMR chemical shifts have been deposited in the BioMagResBank (BMRB: 28091).

## Biostatistics Analysis

All experiments were performed in biological triplicate unless specified. Graphs depict calculated mean values with standard deviation (SD) of triplicates. To assess significance, p-values were determined by student’s t-test and represented as follows: *for p-values < 0.05, ** for p-values < 0.01, *** for p-values < 0.001, and **** for p-values < 0.0001.

## Acknowledgements

This work was supported, in part, by National Institutes of Health Grants R01 GM117437 and R03 CA216134-01, a WW Smith charitable Trust Award, a FCCC/TU Nodal award (XG), R35 GM122517 (RLD) R01NS091336 and R01GM134683 (WP) and a Pre-Pilot Award from U54 CA221704 (ZZ and HF) and funding from NCI CCSG grant P30 CA006927 (XG and RLD).

## Author contributions

X.G. and H.F. wrote the first draft of the manuscript. R.L.D., W.P, A.N.K., R.P., Z.Z and QX edited the manuscript. X.G. and R.L.D. designed experiments. R.P., W.P., and A.N.K., contributed to the experimental design. H.F. performed experiments in Figs. 2-6 and 8, Z.Z. performed experiments in Figs. 1-2, M.A. and A.K. generated constructs and assisted in experiments in Figs. 1-2, FF developed the PP2A/B55α purification scheme for *in vitro* phosphatase experiments, X.W. generated the NMR data and purified proteins for NMR, and Q.X. and R.L.D. performed data-guided computational docking and built a p107 peptide model for B55α binding. XG performed additional analysis of the model.

## Competing interests

The authors declare no potential conflicts of interest.

## Materials & Correspondence

Correspondence to: Xavier Graña, Fels Institute for Cancer Research and Molecular Biology Temple University School of Medicine, AHP Bldg., Room 308 3307 North Broad St. Philadelphia, PA 19140, Tel #: (215) 707-7416, Fax #: (215) 707-2102

## Supplementary Figures

**Supplementary Figure 1.**
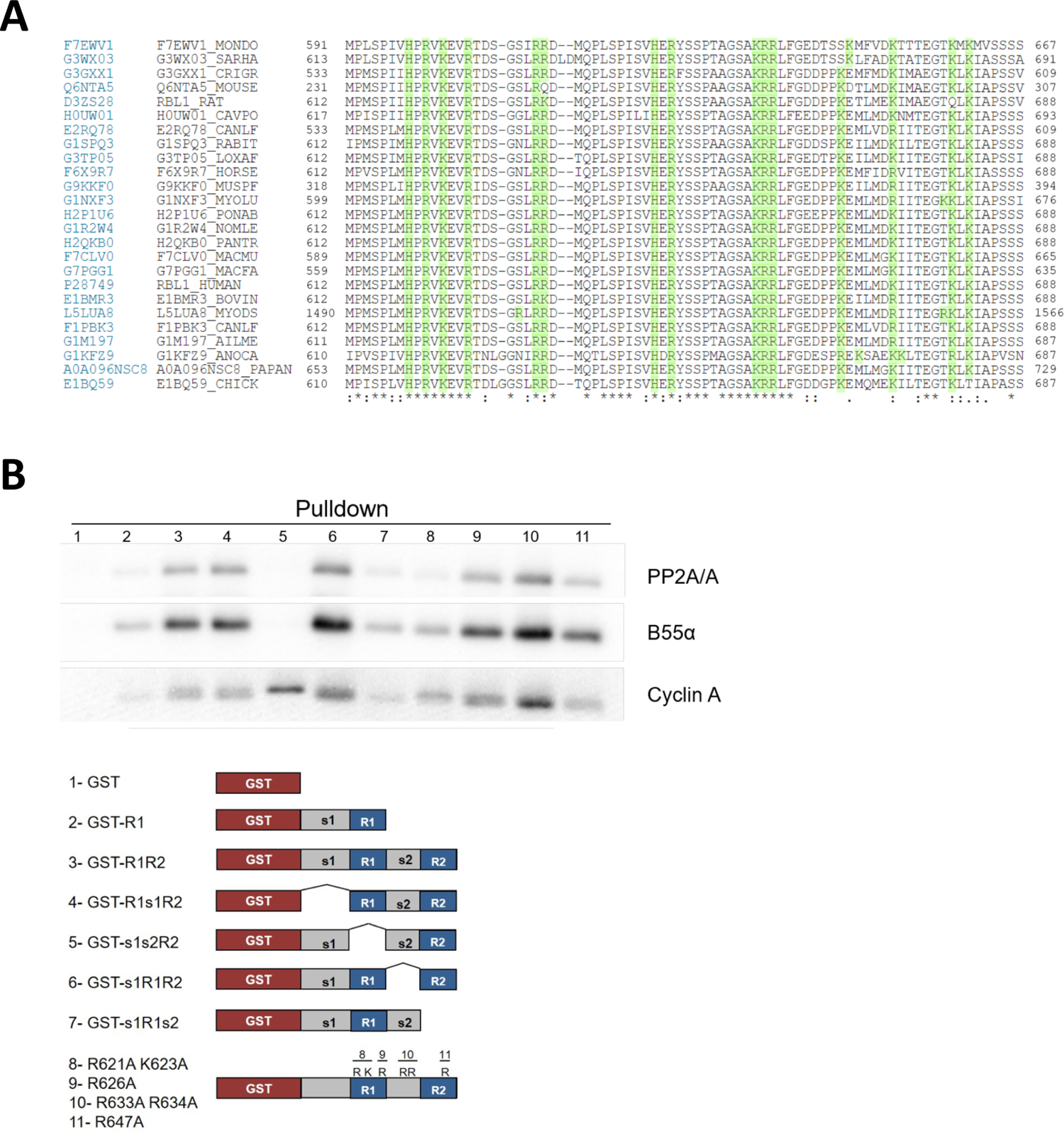
(A) Complete Clustal W alignment of conserved amino acid sequences of the p107 spacer from different species. Three highly conserved regions within the spacer were identified, which are highlighted in grey and named as Region I (R1), Region 2 (R2) and Region 3 (R3). Positively charged residues are highlighted green. (B) Representative pulldown assay of GST-p107 spacer mutants based on the GST-p107 R1R2 construct. Pulldown assays with the indicated fusion proteins from U2-OS lysates were performed and binding of the indicated proteins was determined by western blot analysis.

**Supplementary Figure 2.**
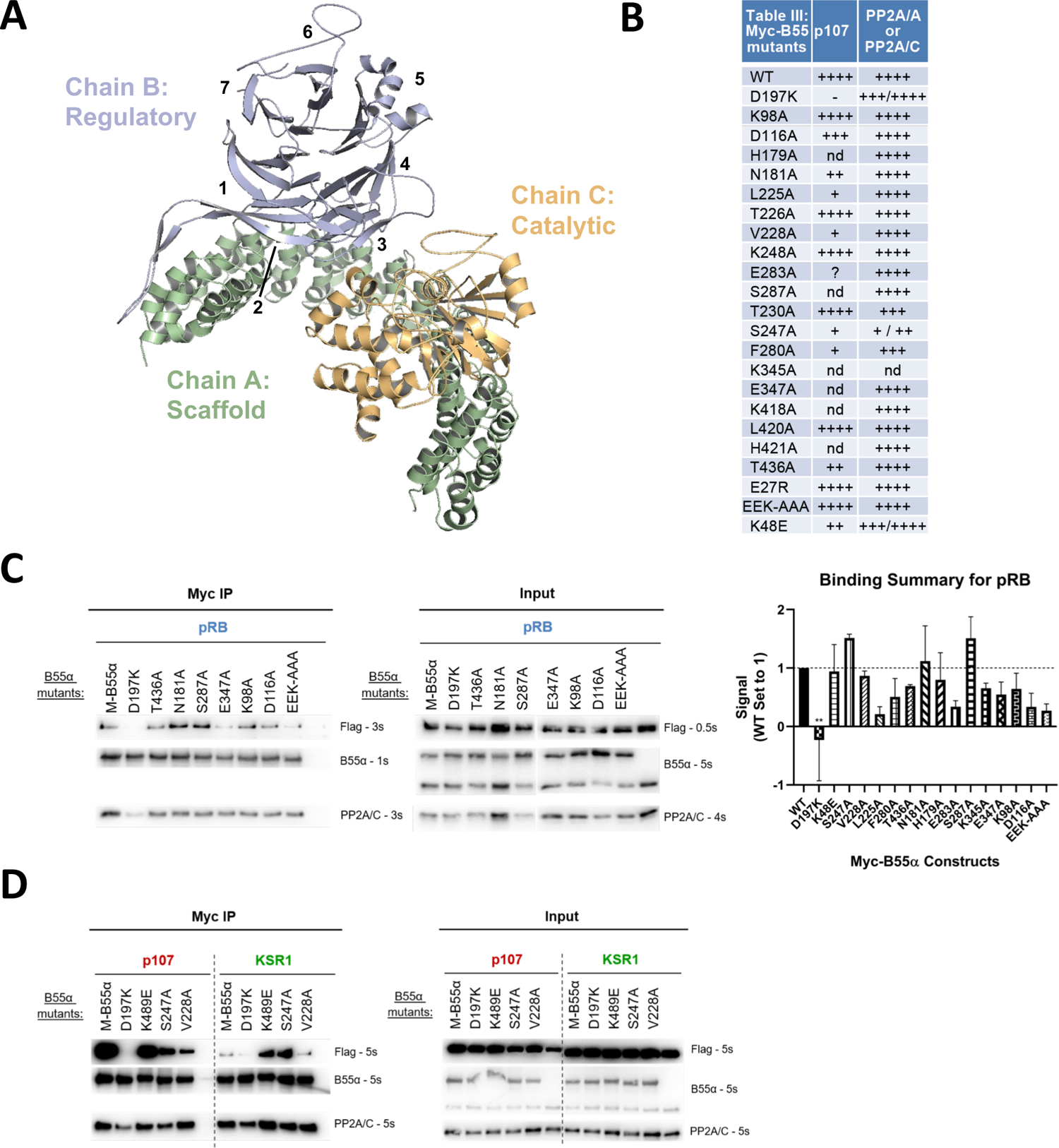
(A) Structure of the PP2A/B55α holoenzyme (PDB: 3DW8). (B) Table depicting complete list of Myc-tagged B55α mutants generated, as well as summarizing the effect on binding to p107 and PP2A/A or PP2A/C based on immunoprecipitation experiments. (C) Representative immunoprecipitation experiment to test pRB binding requirements on Myc-B55α. Flag-pRB and wild-type and mutant Myc-B55α mutant constructs were co-transfected into HEK293T cells and used for IP’s with anti-Myc agarose conjugate. These assays were resolved via SDS-PAGE and proteins were detected using anti-Flag, anti-B55α, and anti-PP2A/C. Mean values for cumulative immunoprecipitation assays for Flag-tagged pRB binding to Myc-B55α constructs are shown right, with statistics indicated above. (D) Representative immunoprecipitation experiment to test KSR1 binding requirements on Myc-B55α. Experiments were performed as above.

**Supplementary Figure 3.**
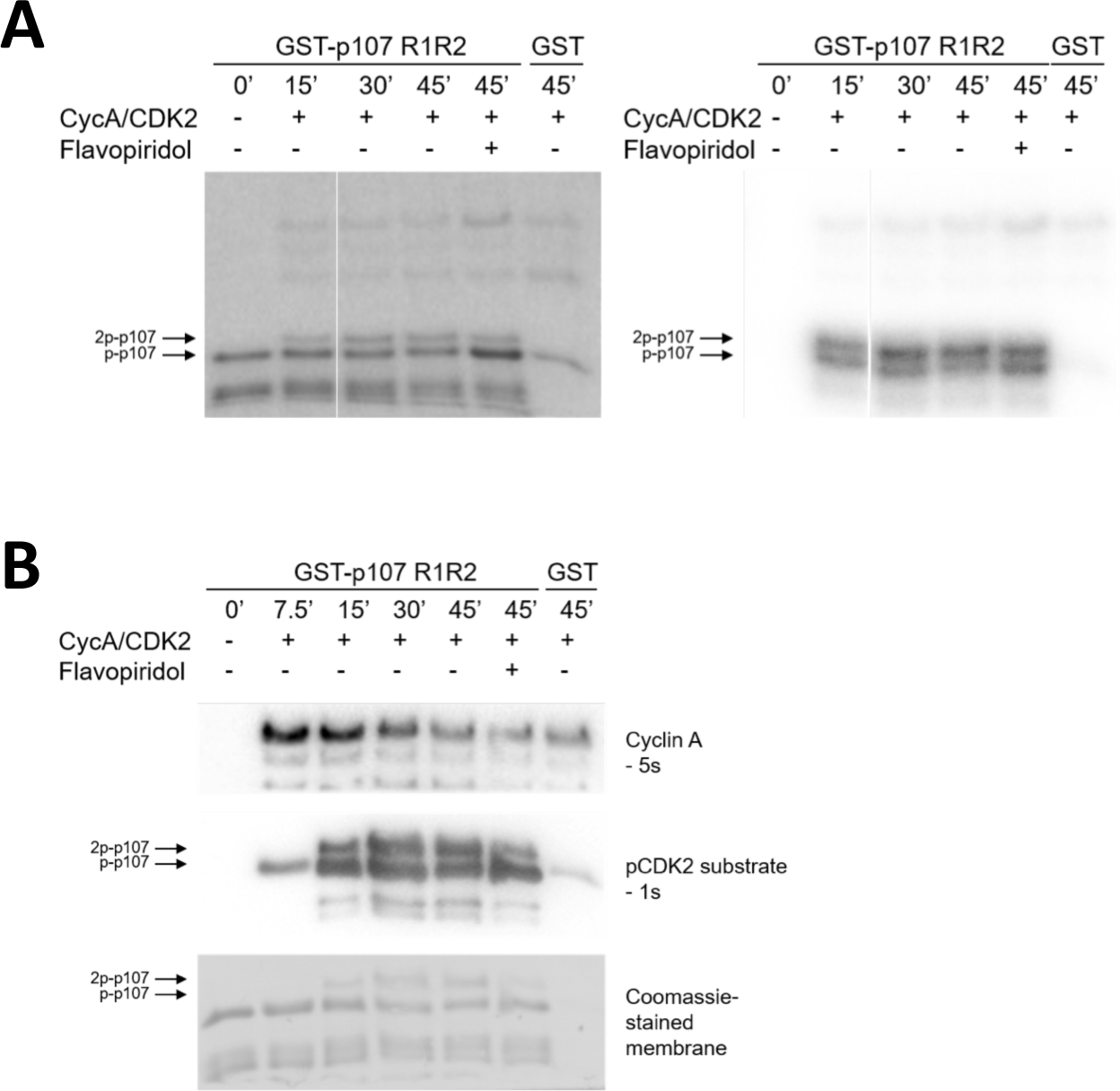
(A) Phosphorylation of purified GST-p107 R1R2 was performed using 0.25 μg recombinant Cyclin A/CDK2 and 5 μCi γ-32P. The indicated time points were collected and samples were resolved via SDS-PAGE. Proteins were detected via Coomassie gel staining and 15 second exposure to a Phosphorimager screen, respectively. (B) Phosphorylation of purified GST-p107 R1R2 was performed using 0.25 μg recombinant Cyclin A/CDK2 and 100 μM ATP. The indicated time points were collected and samples were resolved via SDS-PAGE. Proteins were detected by with anti-cyclin A and anti-pCDK substrate [(K/H)pSP], as well as Coomassie gel staining.

**Supplementary Figure 4.**
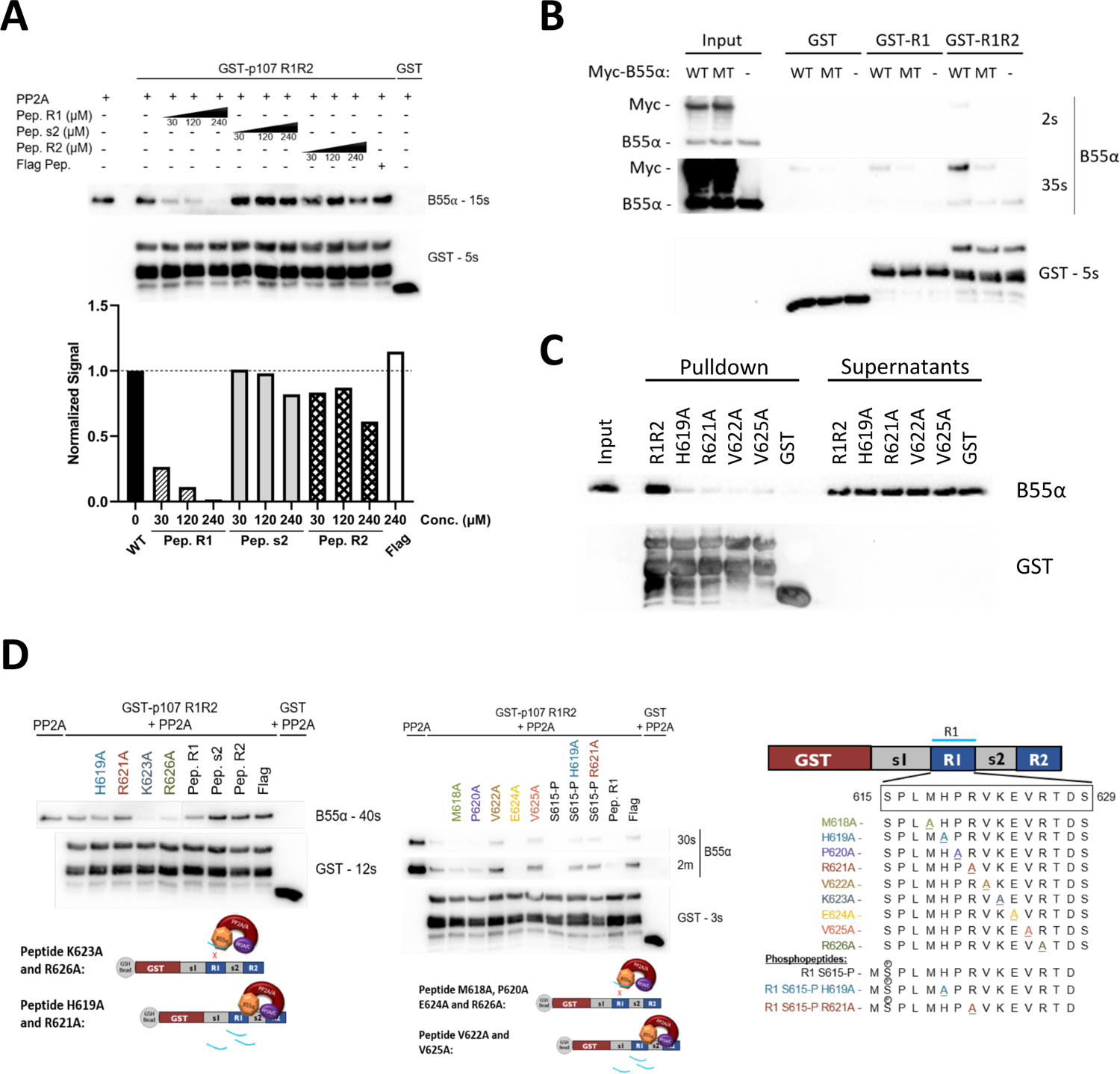
(A) 150 µg of HEK293T lysates was pre-incubated with synthetic p107 peptides and then used in pulldown assays with GST-p107 R1R2 constructs. Proteins were resolved via SDS-PAGE and detected via western blotting using anti-B55α and GST antibodies. Quantification of B55α pulled down relative to the wildtype pulldown was performed for this assay using ImageJ. (B) Myc-B55α wildtype and B55α-D197K mutant constructs were transfected into HEK293T cells and used in pull down assays using the indicated fusion proteins. Samples were resolved via SDS-PAGE and proteins were detected using anti-B55α and anti-GST antibodies. (C) Pulldown assay using GST-R1R2 mutants substituting critical SLiM residues for Ala. Input shown is ∼3% of total volume of PP2A/B55α eluate used in pulldowns. (D) Pulldowns were performed using purified PP2A/B55α and GST-p107 R1R2 with pre-incubations using mutant p107-derived synthetic peptides as well as wildtype phosphopeptides. Proteins were resolved via SDS-PAGE and detected via western blotting using anti-B55α and GST antibodies.

**Supplementary Figure 5.**
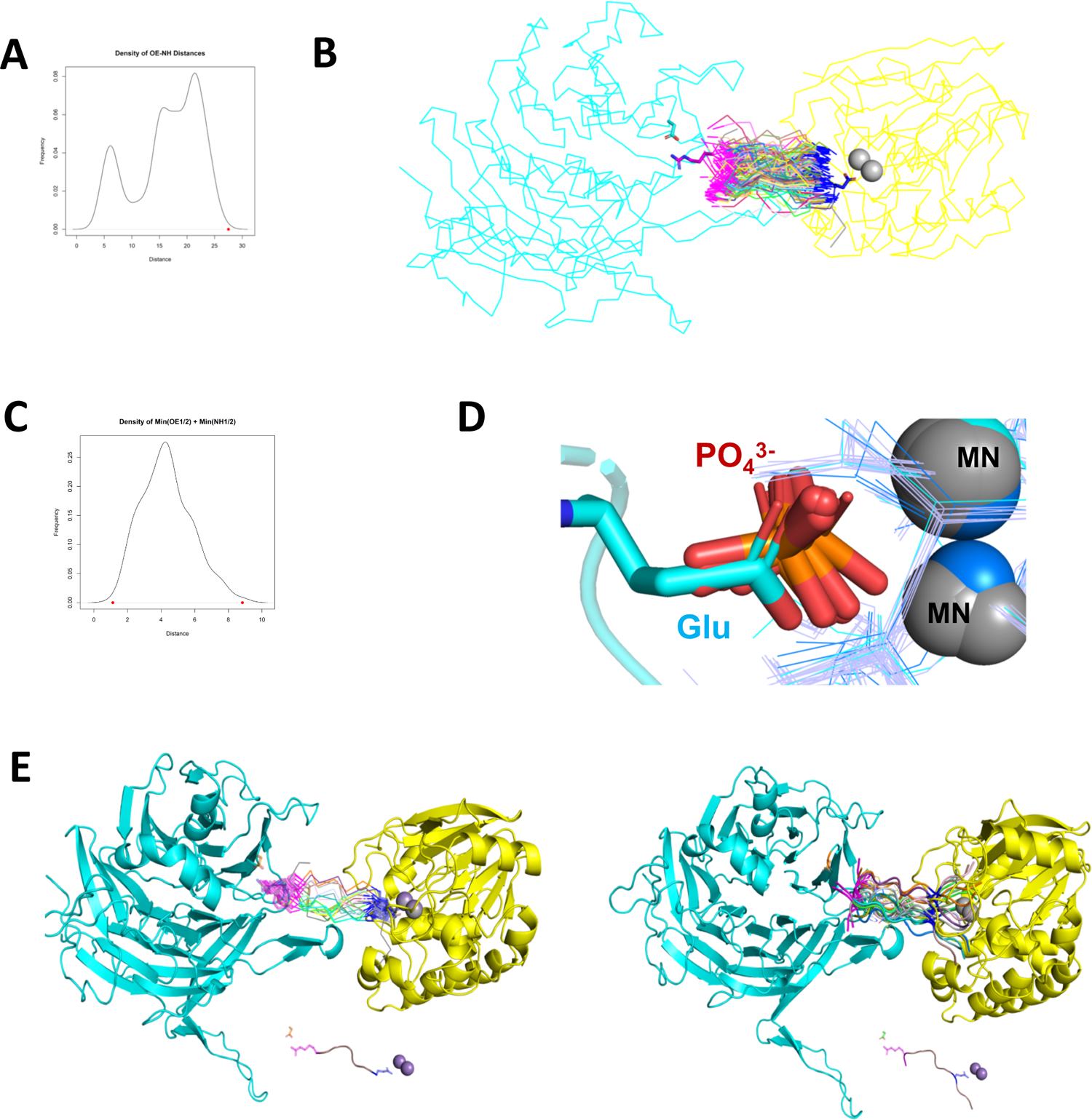
(A) Density plot of distances between OE1/2 of Glu and NH1/2 of Arg in 520 peptide structures with the consensus sequence EPXXXPR. The red dot is the distance between OE1/2 of Glu and NH1/2 of Arg from two reference fragments. (B) Pair fit OE1/2 of Glu residues and NH1/2 of Arg residues of 217 peptide structures to OE1/2 of Glu in MPMEPL fragment and NH1/2 of Arg residue in HPRV fragment. Asp197 is colored in cyan and shown in sticks. Arg in HPRV fragment is colored in magenta and shown in sticks. Glu in MPMEPL fragment is colored in blue and shown in sticks. Peptides with EPXXXPR consensus sequence are shown in ribbon, with their Glu residues are colored in blue and Arg residues are colored in magenta. (C) Density plot of sum of minimum distances (OE1/2 of Glu residues and NH1/2 Arg residues) between each peptide and two reference fragments. The red points are the minimum and maximum values (1.123294 and 8.840146, respectively). (D) Superposing human PPP catalytic subunit structures to PP2A enzyme and the chain of 5HPE, OE1/OE2 atoms of Glu residue are in same positions as oxygen atoms from phosphate groups. (E) (Right) The top 20 peptide structures were aligned to the reference fragments by pair fitting OE1/2 of Glu residues and NH1/2 of Arg residues in PyMOL. (Left) The p107 peptide substrate was built in PyMOL by mutating residues XXX to LMH, and adding MPM residues before the EP and V residue after PR.

**Supplementary Figure 6.**
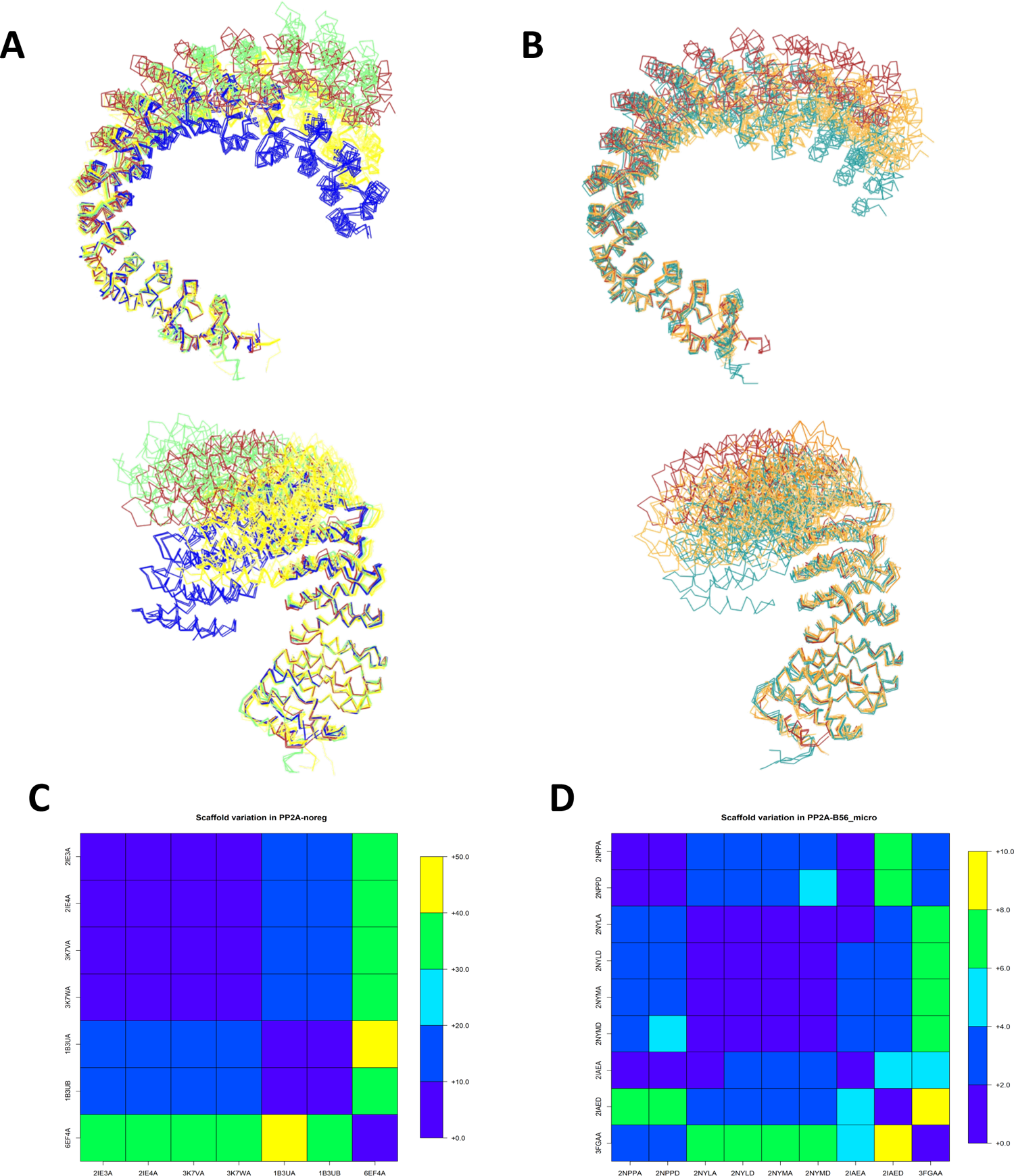
(A) Comparison of the flexibility of the PP2A/A scaffold upon binding to the catalytic subunit and B subunits of the four distinct holoenzymes. B55/B is colored red, B56/B’ is colored yellow, PR70/B‘’ is colored lime, and STRN3/B’’’ is colored blue. (B) Comparison of the flexibility of the PP2A/A scaffold upon binding to TIPRL (colored teal) and small-T antigen (colored orange). B55/B (red) is shown for reference. (C-D) Average distance over 25 residues of scaffold protein in PDB:3DW8 which are in contact with the PP2A enzyme (PDB:3DW8 chain A as reference). (C) PP2A/A structures with no regulatory subunits (PP2A/A noreg). 6EF4A corresponds to PP2A/A scaffold without catalytic subunit. (D) Scaffold structures with B56 and microcystin (PP2A-B56_micro).

